# Integrated annotations and analyses of small RNA-producing loci from 47 diverse plants

**DOI:** 10.1101/756858

**Authors:** Alice Lunardon, Nathan R. Johnson, Emily Hagerott, Tamia Phifer, Seth Polydore, Ceyda Coruh, Michael J. Axtell

## Abstract

Plant endogenous small RNAs (sRNAs) are important regulators of gene expression. There are two broad categories of plant sRNAs: microRNAs (miRNAs) and endogenous short interfering RNAs (siRNAs). MicroRNA loci are relatively well-annotated but comprise only a small minority of the total sRNA pool; siRNA locus annotations have lagged far behind. Here, we used a large dataset of published and newly generated sRNA sequencing data (1,333 sRNA-seq libraries containing over 20 billion reads) and a uniform bioinformatic pipeline to produce comprehensive sRNA locus annotations of 47 diverse plants, yielding over 2.7 million sRNA loci. The two most numerous classes of siRNA loci produced mainly 24 nucleotide and 21 nucleotide siRNAs, respectively. 24 nucleotide-dominated siRNA loci usually occurred in intergenic regions, especially at the 5’-flanking regions of protein-coding genes. In contrast, 21 nucleotide-dominated siRNA loci were most often derived from double-stranded RNA precursors copied from spliced mRNAs. Genic 21 nucleotide-dominated loci were especially common from disease resistance genes, including from a large number of monocots. Individual siRNA sequences of all types showed very little conservation across species, while mature miRNAs were more likely to be conserved. We developed a web server where our data and several search and analysis tools are freely accessible at http://plantsmallrnagenes.science.psu.edu.

## Introduction

Plant regulatory small RNAs (sRNAs) play important roles in almost all biological processes. Endogenous sRNAs are 20-24 nucleotides in length and derive from longer RNA precursors that are processed by DICER-LIKE (DCL) ribonucleases. Once processed, they are loaded into Argonaute (AGO) proteins to form the RNA-induced silencing complex (RISC). Then, sRNAs guide the RISC complex to complementary sites on target RNAs, inducing either post-transcriptional or transcriptional gene silencing.

Endogenous sRNAs can be grouped in two broad classes based on their biogenesis and typical functions: microRNAs (miRNAs) and small interfering RNAs (siRNAs) (Axtell 2013a). MiRNAs are typically 21-22 nucleotides long, processed from single-stranded RNA (ssRNA) stem-loop precursors by DCL1, and regulate gene expression post-transcriptionally, directing mRNA degradation and translational repression (Rogers and Chen 2013). SiRNAs are processed from double-stranded RNA (dsRNA) precursors and are categorized in multiple sub-classes. The most abundant sub-class of siRNAs participates in the RNA-directed DNA methylation (RdDM) pathway, involving 24 or 21-22 nucleotide siRNAs. 24 nucleotide siRNAs are derived from Polymerase IV (Pol IV) transcripts that are converted to dsRNAs by RNA-dependent RNA polymerase 2 (RDR2) which are then processed by DCL3. They act in “canonical” RdDM, primarily targeting transposable elements (TEs) and other repeats to induce DNA methylation and reinforce transcriptional silencing. 21-22 nucleotide siRNAs are derived from Pol II transcripts and are copied by RDR6 into dsRNAs and processed by DCL2/DCL4. They act in the non-canonical RdDM pathway to establish the silencing of young TEs, both transcriptionally and post-transcriptionally (Nuthikattu et al. 2013). Another major siRNA sub-class is secondary siRNAs. Their biogenesis is triggered by a miRNA-directed cleavage of a coding or non-coding transcript. The transcript is then converted to dsRNA by RDR6 and processed by DCL proteins into secondary siRNAs in a phased pattern relative to the miRNA cut site. Phased secondary siRNAs (phasiRNAs) are typically 21 or 22 nucleotides long, however a specific population of 24 nucleotide phasiRNAs has been detected in anthers of many angiosperms (Xia et al. 2019). *TAS* genes are an example of loci generating non-coding RNA precursors that produce secondary siRNAs, which act *in trans* (trans-acting siRNAs, tasiRNAs) on other targets and direct their cleavage (Allen et al. 2005). Pentatricopeptide repeat (PPR) genes are the first reported protein-coding genes generating secondary siRNAs in *Arabidopsis thaliana* (Howell et al. 2007).

At the chromosomal level, sRNA distribution correlates with gene density, typically lower in the centromeric and pericentromeric regions and enriched in the distal euchromatic regions. This trend has been observed in maize (He et al. 2013), rice (Wei et al. 2014), tomato (The Tomato Genome Consortium 2012), hot pepper (Kim et al. 2014), upland cotton (Song et al. 2015) and sugar beet (Dohm et al. 2014). However, in a smaller number of species sRNAs mostly arise from centromeric and pericentromeric regions away from genes, as shown in *A. thaliana* (Kasschau et al. 2007; Ha et al. 2009), soybean (Schmitz et al. 2013), cucumber (Lai et al. 2017) and *Brachypodium distachyon* (The International Brachypodium Initiative 2010).

Despite differences in chromosomal distributions, the sRNA profiles near protein-coding genes are conserved amongst plant species, with 24 nucleotide siRNAs preferentially found in gene-proximal regions but depleted in gene bodies themselves. This pattern has been described in maize (Gent et al. 2013), rice (Wei et al. 2014), rapeseed (Shen et al. 2017), Chinese cabbage (Woodhouse et al. 2014), soybean (Song et al. 2013), upland cotton (Song et al. 2015) and *A. thaliana* (Kasschau et al. 2007; Ha et al. 2009). Depending on the species, sRNAs have opposite effects on the regulation of the proximal downstream genes. In maize, 24 nucleotide siRNAs are found with higher probability near expressed genes than non-expressed genes (Gent et al. 2013; Lunardon et al. 2016). Here, siRNAs participate in RdDM to reinforce the silencing of TEs that are inserted upstream of genes, where the chromatin is accessible, therefore repressing the potentially deleterious Pol II transcription of TEs (Gent et al. 2014). In contrast, the siRNA-mediated silencing of TEs near genes is linked to lower expression of the genes in *A. thaliana* and Chinese cabbage (Hollister et al. 2011; Woodhouse et al. 2014). In addition to target TEs near genes, 24 nucleotide siRNAs can also target TEs inserted inside genes, affecting their expression (Wei et al. 2014; Lunardon et al. 2016).

Genome-wide analyses in barley, soybean, *Medicago truncatula* and *Physcomitrella patens* showed that 21 nucleotide siRNAs are not enriched in gene body regions (Hackenberg et al. 2016; Schmitz et al. 2013; Lelandais-Brière et al. 2009; Coruh et al. 2015). Nevertheless, there are many cases of well characterized genes generating 21 nucleotide phasiRNAs in dicots: nucleotide binding/leucine-rich repeat (NB-LRR) and receptor like kinase (RLK) resistance genes, PPR genes, auxin-responsive factor (ARF) genes, MYB and NAC transcription factors and F-BOX genes (Arikit et al. 2014; Hu et al. 2015a; Xia et al. 2015b). NB-LRR genes evolve rapidly by tandem duplication and they are controlled by sRNA-mediated silencing to avoid their over-expression and prevent autoimmune responses (Yang and Huang 2014). This mechanism is conserved in a large number of dicots: soybean, *M. truncatula*, common bean, chickpea, *Populus trichocarpa*, cassava, pima cotton, potato and Norway spruce (Zhai et al. 2011; Formey et al. 2015; Srivastava et al. 2015; Klevebring et al. 2009; Xia et al. 2014; Hu et al. 2015b; Xia et al. 2015a). Amongst monocots, 21 nucleotide phasiRNAs from NB-LRR genes have been only found in barley and wheat so far (Liu et al. 2014; Zhang et al. 2019). This is consistent with the fact that monocots, in contrast to dicots, produce phasiRNAs mainly from non-coding RNAs (Komiya 2017; Zheng et al. 2015).

The conservation of sRNAs across plants has been widely investigated for miRNAs. There are deeply conserved miRNA families together with their targets, suggesting common functional regulatory networks (Axtell and Bowman 2008). However, the majority of miRNA sequences are species-specific, indicating the presence of numerous young or still evolving miRNAs (Cuperus et al. 2011; Chávez Montes et al. 2014). Much less is known about conservation of siRNAs but a study comparing *Arabidopsis thaliana* and *Arabidopsis lyrata* suggested that individual siRNA sequences are not conserved even between closely related species (Ma et al. 2010). Moreover, while in most species analyzed so far the 24 nucleotide siRNAs are the most abundant expressed group of sRNAs, mosses, lycophytes and conifers lack a strong peak of 24 nucleotide siRNAs (Axtell and Bartel 2005; Banks et al. 2011; Dolgosheina et al. 2008).

There are several existing web-based resources that serve sRNA sequencing (sRNA-seq) data for multiple plants. The Cereal small RNA Database contains maize and rice genome browsers with accessible sRNA-seq data (Johnson et al. 2007). The Pln24NT website stores annotations and sequences of 24 nucleotide siRNA reads and loci for 10 species (Liu et al. 2017). The Next-Gen Sequence Databases produced by the Meyers lab contain sRNA-seq and other high-throughput data with custom-built genome browsers and search functions for 27 species (Nakano et al. 2006). The miRBase database (Kozomara and Griffiths-Jones 2014) provides curated, comprehensive annotations of *MIRNA* loci in a very large number of species. An equivalent database for the storage and distribution of reference annotations of siRNA-producing loci in a vast number of plant genomes does not exist (Coruh et al. 2014).

In this study, we used a large dataset of published and newly generated sRNA-seq data, that we processed with a consistent pipeline, to create reference sRNA loci annotations for 47 plant species, including model plants and crops. We propose and use a systematic nomenclature and ontology for sRNA-producing loci that is consistent with their biology and easily traceable and updatable. We examined the genome-wide distribution of sRNA loci relative to protein-coding genes and compared it across species, providing insights into conserved sRNA functions. We organized the sRNA-seq alignment data and sRNA loci annotations in a freely available web-based database that represents an important public resource for future studies aimed to understand the biological function of sRNAs.

## Results

### Identification and classification of sRNA loci in 47 plants

We obtained and analyzed 48 plant genome assemblies, representing 47 different species (Table 1; two independent assemblies of *Cuscuta campestris* were analyzed). To facilitate succinct communication in figures and our database, a short code was designated for each assembly. The code begins with a three-letter prefix representing the genus and species, following the abbreviations established by miRBase (Kozomara et al. 2019). The second part of the code indicates the genome build (’-b’) version in use. These genome assemblies varied widely in size, contiguity, protein-coding gene number, and repeat content (Supplemental Fig. S1,S2). Most genome assemblies were from crops; others included the model plants *Arabidopsis thaliana* and *Medicago truncatula*, the parasitic plant *Cuscuta campestris*, and representatives of diverse lineages (*Amborella trichopoda* [basal angiosperm], *Picea abies* [gymnosperm], *Physcomitrella patens* [bryophyte], and *Marchantia polymorpha* [bryophyte]).

**Table 1.**
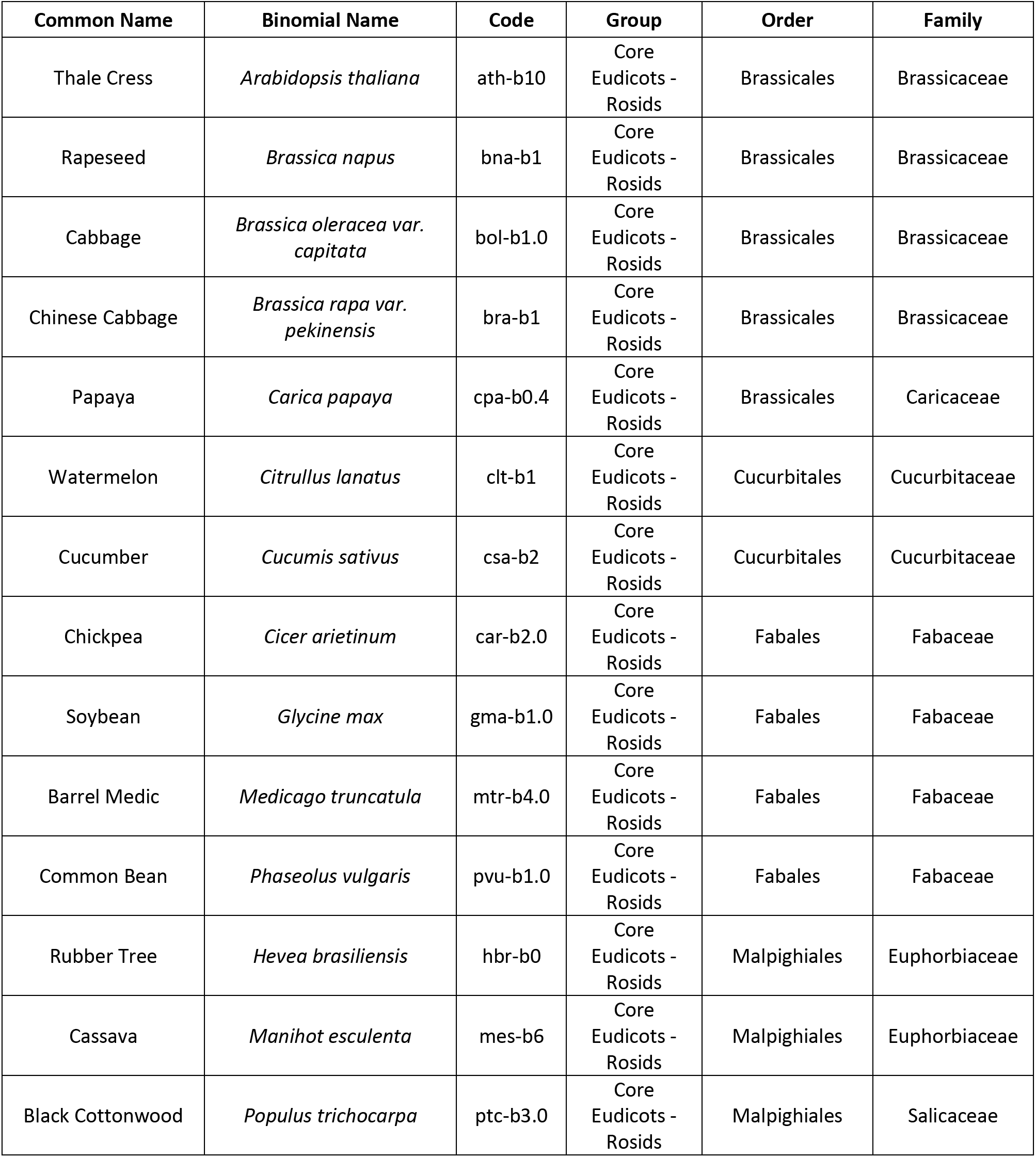
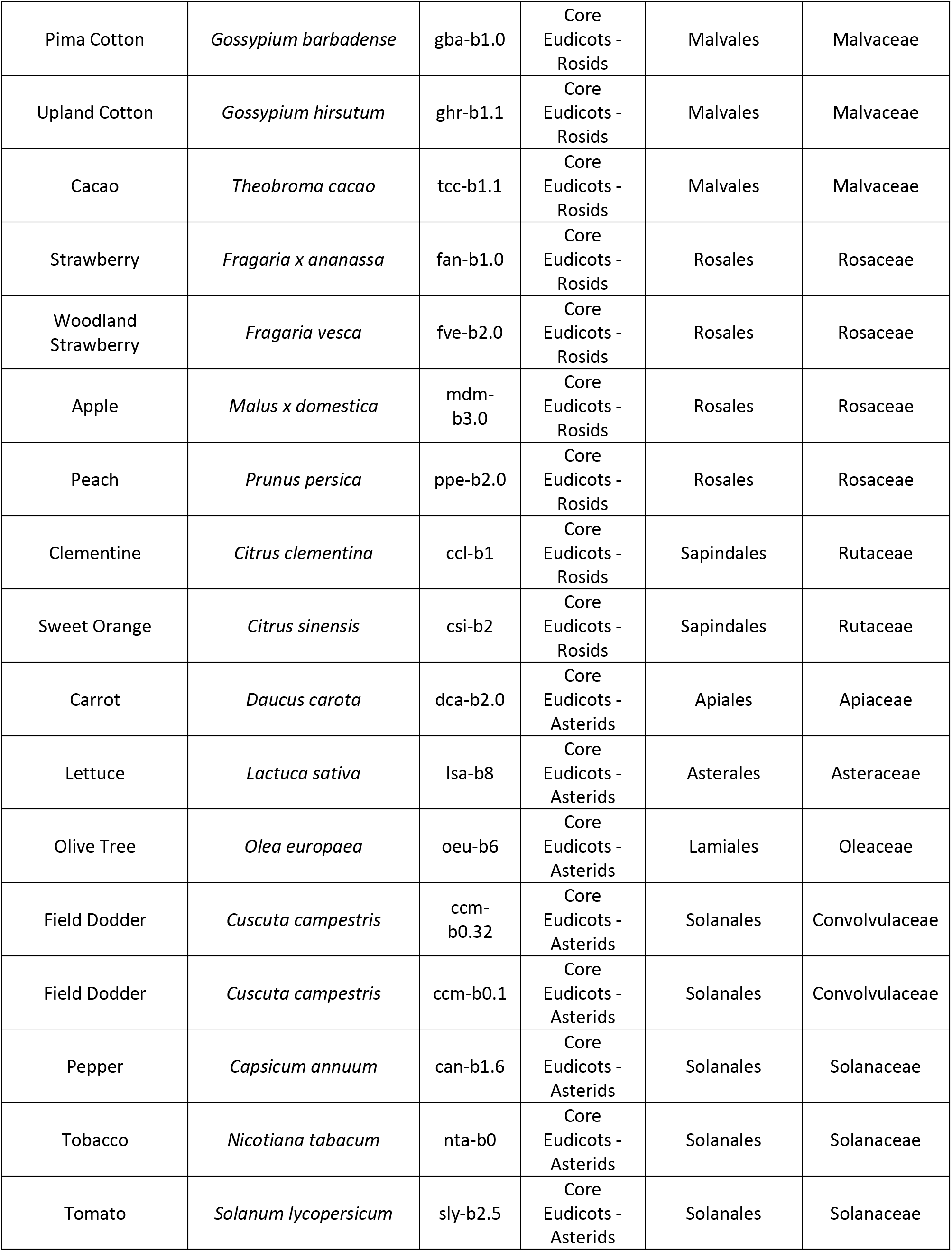
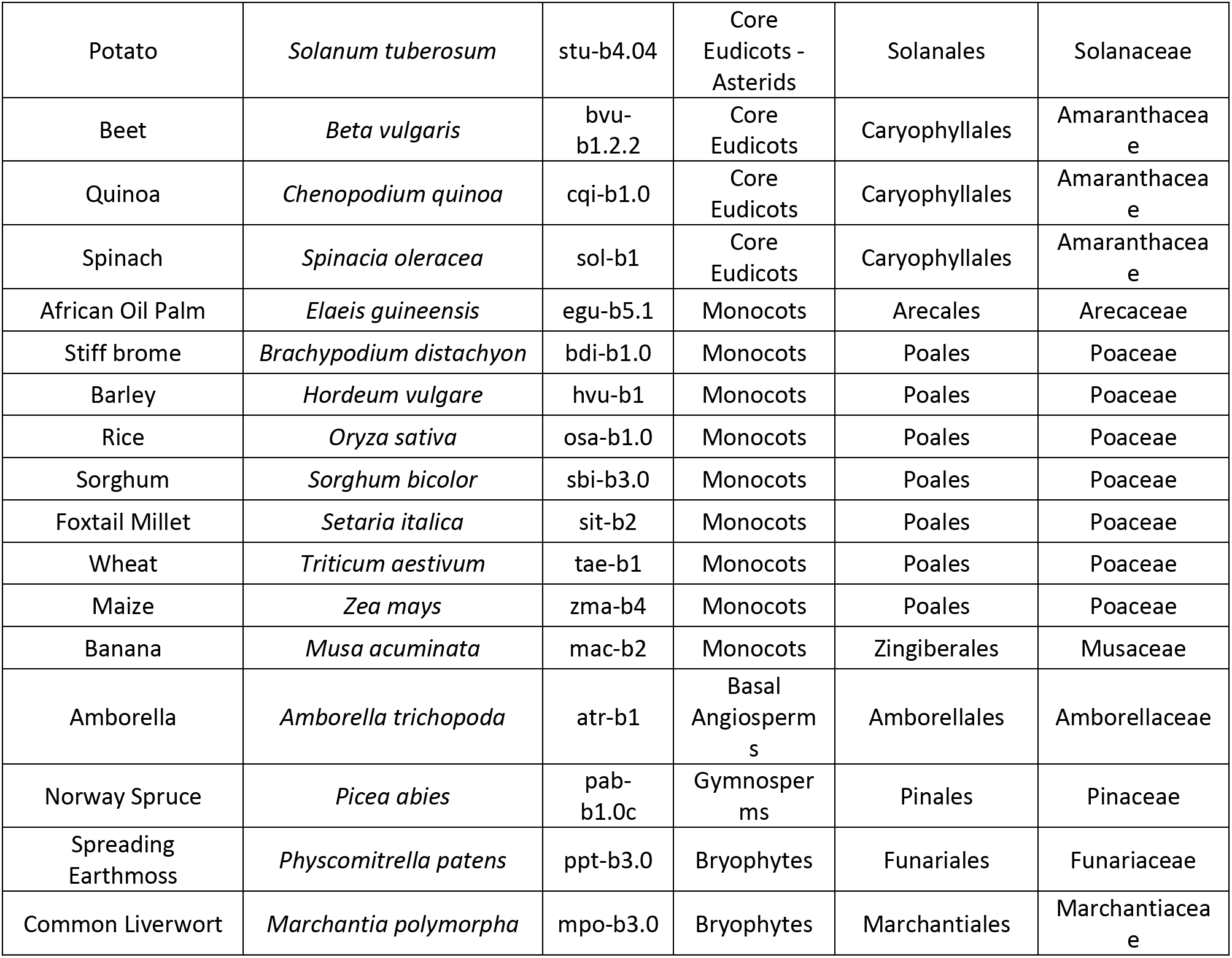
Plants included in this study.

We gathered sRNA-seq libraries from each genome (Figure 1A). In most cases, these data were from public sequencing archives (Supplemental Table S1). In a few cases, we also generated novel sRNA-seq libraries (*Zea mays*, *Spinacia oleracea*, *Daucus carota, Theobroma cacao*; Supplemental Table S1). We sought to annotate the full diversity of sRNA loci and thus selected libraries with the goal of including as many different tissues and conditions as possible. However, we excluded low-depth sRNA-seq datasets (less than two million reads aligned to the genome) and also excluded sRNA-seq datasets from mutants known to affect sRNA biogenesis or stability. For each given genome assembly, all cognate sRNA-seq libraries were aligned and then merged into a single master sRNA alignment which we call the “reference set” (Figure 1A). Reference sets had considerable variation in both total number of sRNA reads (minimum: 2.1E6, median: 1.6E8, maximum: 4.1E9) and in number of contributing sRNA-seq libraries (minimum: 1, median: 11, maximum: 161)(Supplemental Fig. S3).

**Figure 1.**
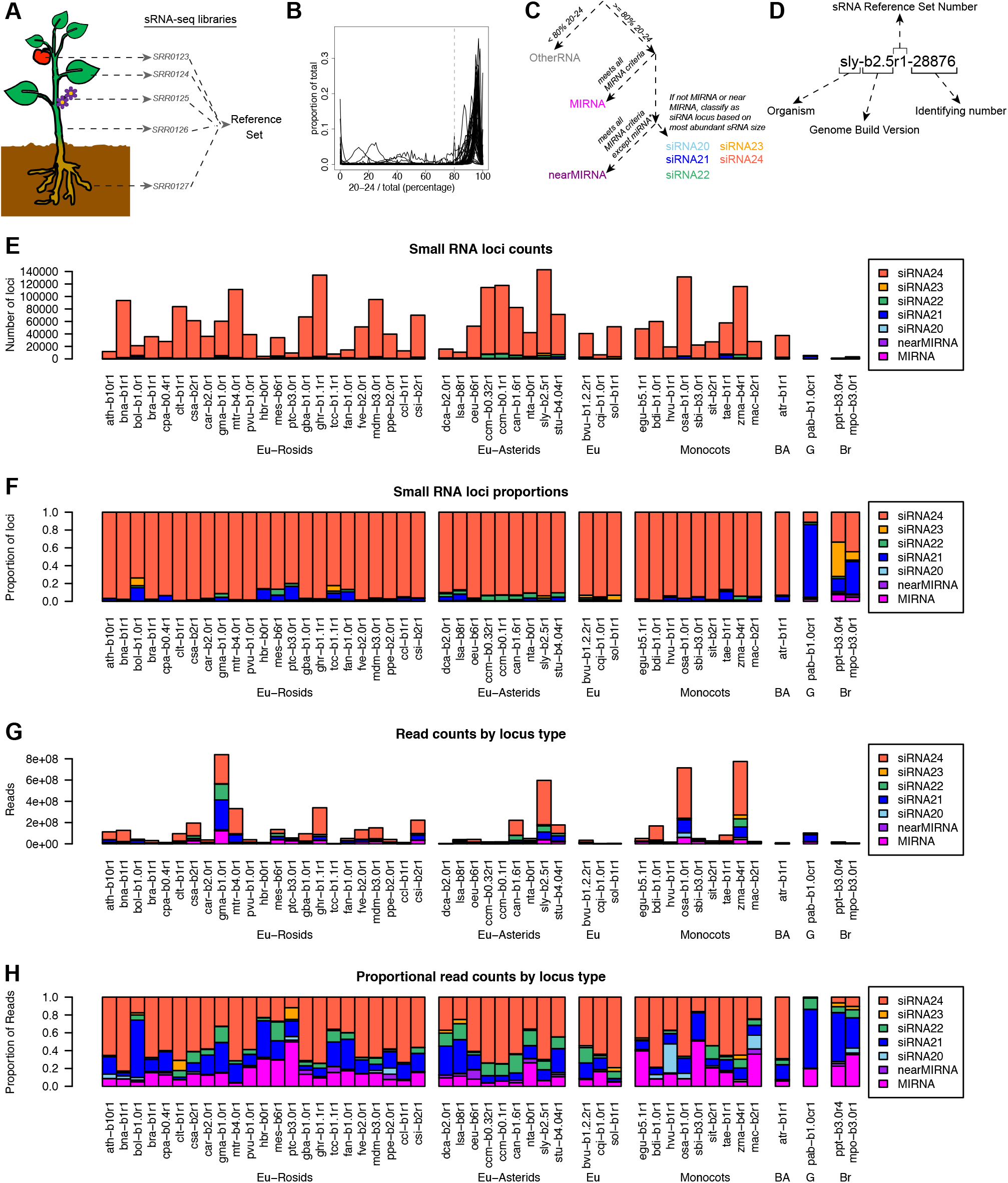
Overview of sRNA locus annotation pipeline and summary of annotated sRNA loci. **(A)** Schematic illustrating how multiple sRNA libraries from diverse plant tissues are merged to create a ‘reference set’ of sRNAs for a given species. Accession numbers shown are fictional. **(B)** Distributions of the fractions of sRNAs between 20-24 nucleotides in length (inclusive) within all loci in each genome. Gray line at 80% represents the cutoff used to discriminate silencing-related RNA loci from other types of sRNA-producing loci. **(C)** Flowchart illustrating the ontology used to classify sRNA-producing loci. Colors designating different locus types are used throughout this work. **(D)** Schematic illustrating the nomenclature used to annotate sRNA-producing loci. **(E-H)** Summary of annotated sRNA loci, by species and locus type, excluding the category ‘OtherRNA’ **(E)** Counts of annotated loci. **(F)** Proportions of annotated loci. **(G)** Total counts of aligned small RNAs in reference sets. **(H)** Proportions of small RNA total read counts in reference sets. See Table 1 for species codes. Eu: eudicots, BA: basal angiosperm, G: gymnosperm, Br: bryophyte.

For annotation, we first identified genomic regions producing sRNAs, independently in all sRNA-seq libraries with ShortStack (Axtell 2013b; Johnson et al. 2016). Then we compared the sRNA expression from different samples of the same species and identified the regions that were robustly expressing sRNAs in at least three separate samples. Millions of discrete sRNA clusters were annotated in this way and defined as sRNA-producing loci, which were then analyzed in the genome-aligned reference sets. Canonical plant miRNAs and siRNAs are between 20 and 24 nucleotides in length, while other types of sRNA loci produce a broader range of RNA sizes. For each locus, we computed the fraction of aligned sRNA-seq reads that were 20-24 nucleotides long. We found that these fractions had consistent bimodal distributions in the various genomes (Figure 1B). Based on these distributions, we used a cutoff of 80% to discriminate canonical siRNA/*MIRNA* loci from ‘OtherRNA’ loci (Figure 1C). We then developed a simplified ontology to describe the siRNA and *MIRNA* loci: ‘MIRNA’ loci were those that met all *MIRNA* annotation criteria, while ‘nearMIRNA’ loci met most criteria except for that the exact predicted miRNA*, the complementary strand to the mature miRNA in the miRNA-miRNA* duplex, was not sequenced. The remaining loci were classified as siRNA loci based on the predominant length of aligned sRNAs within each locus (Figure 1C). This ontology has the advantage of being applicable to any genome regardless of any other annotations or information. We also devised a simple nomenclature to systematically name the sRNA loci (Figure 1D). In total, we annotated approximately 2.7E6 sRNA-producing loci from the 48 genome assemblies (Supplemental Table S2; also see http://plantsmallrnagenes.science.psu.edu for easier access and more analysis options).

The ‘OtherRNA’ category of loci, defined by having less than 80% of aligned reads with sizes between 20-24 nucleotides in length, typically comprised less than half of all loci in the flowering plants (Supplemental Fig. S4A,B). In contrast, the majority of loci identified in one gymnosperm and two bryophyte genomes were annotated as OtherRNA (Supplemental Fig. S4A,B). Across all taxa, OtherRNA loci typically contributed large fractions of total read abundance (Supplemental Fig. S4C,D). This is because many of the OtherRNA loci represented clusters of short fragments derived from highly abundant, longer RNAs, such as rRNAs, tRNAs, and plastid-derived mRNAs. There is evidence that some plant RNAs longer than 24 nucleotides, or shorter than 20 nucleotides, may function as gene-regulatory factors (Martinez et al. 2017); such loci will have been annotated in the OtherRNA category by our procedure. Nonetheless, we focused our subsequent analyses on the MIRNA, nearMIRNA, and siRNA loci dominated by 20-24 nucleotide RNAs because these sizes are most clearly associated with production by DCL endonucleases and usage by AGO proteins. By default, ShortStack assigns a phasing score to the sRNA loci based on the algorithm described in Guo et al. 2015. However, an accurate annotation of the phasing would require a more complex study to avoid false positives that may be produced by the commonly used phasing-detecting algorithms (Polydore et al. 2018). Therefore, we did not further analyze the phasing of the sRNA loci in this analysis.

After excluding OtherRNA loci, the remaining loci were mostly designated siRNA24 in angiosperms (Figure 1E-F). In contrast, and consistent with prior reports (Dolgosheina et al. 2008; Axtell and Bartel 2005), gymnosperm and bryophyte loci were less dominated by the siRNA24 type and instead had more siRNA21 loci. When tallied by sRNA abundance, MIRNA and siRNA21 loci made substantial contributions in all taxa (Figure 1G-H). This indicates that a relatively small number of MIRNA and siRNA21 loci produce high levels of their respective sRNAs. In a number of species, the proportion of 22 nucleotide siRNAs was also substantial and this trend was particularly consistent amongst the asterids (Figure 1H). In most cases, angiosperms had more annotated sRNA loci compared to non-angiosperms (Supplemental Fig. S4A, Figure 1E). However, that comparison is potentially complicated by the different amounts of input sRNA reads used for each species (Supplemental Fig. S3).

### The plantsmallrnagenes.science.psu.edu server

All data and analyses from this study have been systematically organized and are freely available at https://plantsmallrnagenes.science.psu.edu. Users can search for loci of interest by sRNA sequence, *MIRNA* family name, locus name, or by BLAST-based homology searches. A JBrowse-based genome browser is available for each of the 48 genomes. Genome browsers are customized to display sRNA-seq data based on sRNA size, strand, and multi-mapping (Figure 2A). Genome browsers also allow users to highlight a region of interest and perform on the fly analyses, including ShortStack (Axtell 2013b; Johnson et al. 2016) and visualization of possible *MIRNA* hairpins (Figure 2B). Bulk data are also available in standard, widely used formats: sRNA-seq alignments are in the BAM format, while annotations of sRNA loci are in the GFF3 format. It is our intention to maintain and expand this resource for the benefit of anyone interested in the analysis of plant sRNA-producing loci.

**Figure 2.**
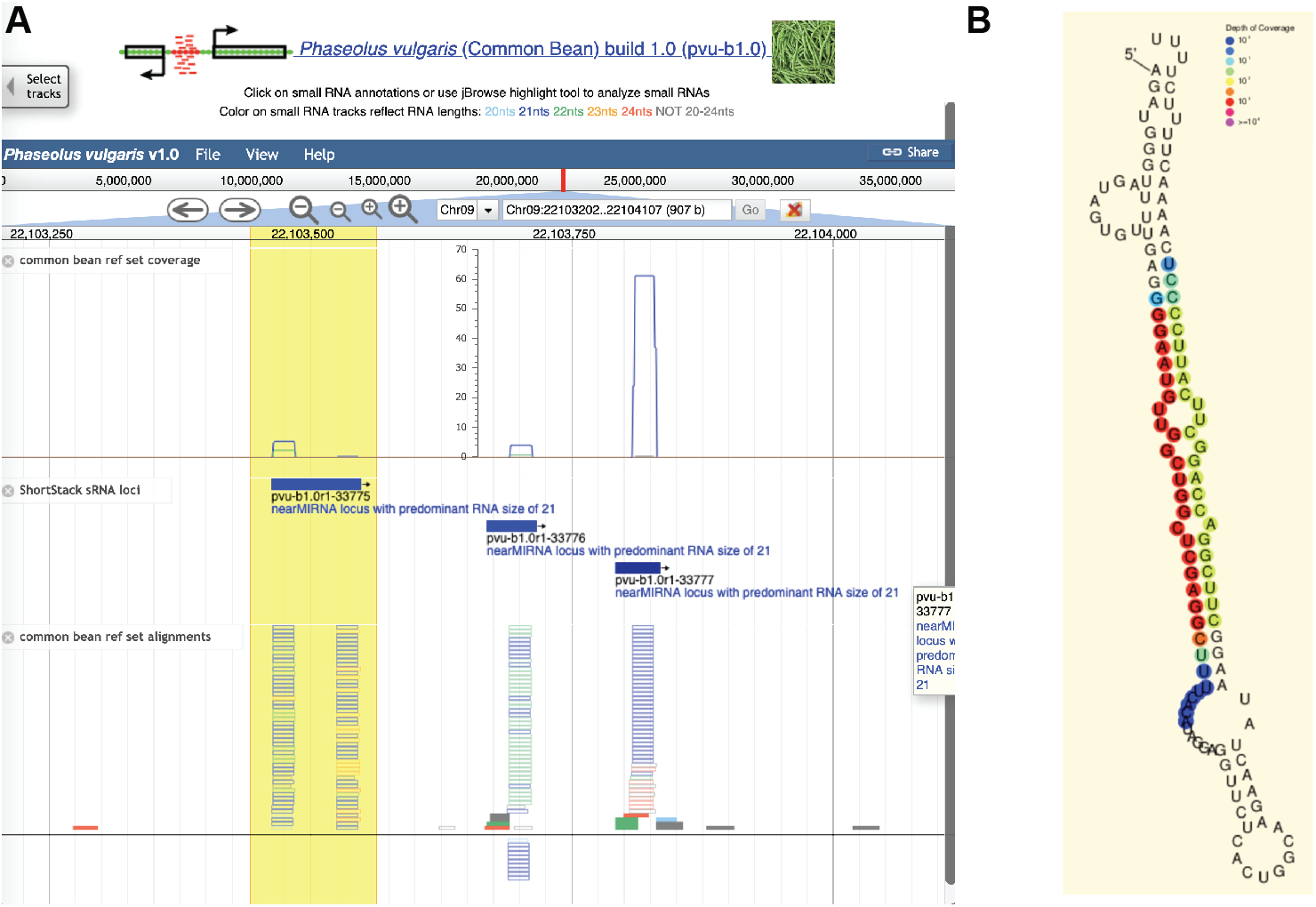
Example screenshots from https://plantsmallrnagenes.science.psu.edu **(A)** Screenshot of genome browser for a region of *Phaseolus vulgaris* chromosome 9. Coverage track shows sRNA-seq alignment depths from the reference set, separated by sRNA lengths (indicated by colors). ShortStack sRNA loci track shows sRNA locus annotations. Alignments track shows individual sRNA reads from the reference set, with lengths indicated by colors. Hollow bars indicate multi-mapped reads; solid bars are uniquely mapped reads. A user-highlighted region is indicated in yellow. **(B)** Analysis of predicted RNA secondary structure with sRNA-alignment depths indicated by colors. This analysis is one of several that can be triggered by user selection of a region of interest (yellow region in panel A).

### Chromosomal distribution of sRNA loci and association with protein-coding genes

Where feasible based on genome assembly quality, we compared the distribution of sRNA loci and genes across entire chromosomes and confirmed that the most common trend is a positive correlation between gene density and sRNA density (Supplemental Fig. S5), as has previously been shown in several prior species-specific studies (He et al. 2013; Wei et al. 2014; The Tomato Genome Consortium 2012; Kim et al. 2014; Song et al. 2015; Dohm et al. 2014). *A. thaliana* is unique in that it has a clear trend from telomeres to centromeres of decreasing gene density and increasing sRNA loci density (Kasschau et al. 2007; Ha et al. 2009). Surprisingly, rice showed a similar trend to *A. thaliana* (Supplemental Fig. S5). Chinese cabbage and sweet orange also showed a slight inverse correlation between the gene and the sRNA loci distributions. Finally, soybean had a general positive correlation between genes and sRNA loci but in the most distal segments of the chromosome arms it showed a local negative correlation (Supplemental Fig. S5).

We examined siRNA21 loci and siRNA24 loci locations relative to protein-coding genes. Other types of sRNA loci were excluded due to their lower frequencies. Coverage of protein-coding genes and flanking 5kb regions by siRNA21 or siRNA24 loci was calculated and normalized. siRNA21 loci had a striking tendency in nearly all taxa to overlap with protein-coding genes (Figure 3A). In contrast, siRNA24 loci were strongly depleted in protein-coding genes in most angiosperms (Figure 3B). siRNA24 loci were often strongly enriched in the 5’-proximal regions upstream of protein-coding genes. There were, however, some notable exceptions to this pattern. There was no upstream peak of siRNA24 loci in bryophytes and the gymnosperm (Figure 3B), which is consistent with the generally low levels of siRNA24 loci in these taxa (Figure 1). The basal angiosperm *Amborella trichopoda* was unusual in that siRNA24 loci were not depleted in gene bodies at all (Figure 3B). Finally, the model plant *A. thaliana* also lacked a conspicuous upstream gene-proximal enrichment of siRNA24 loci. This observation, together with the unique chromosomal distribution of sRNA loci in *A. thaliana*, suggests that *A. thaliana* may not be representative of most angiosperms in its genome-wide patterns of sRNA loci.

**Figure 3.**
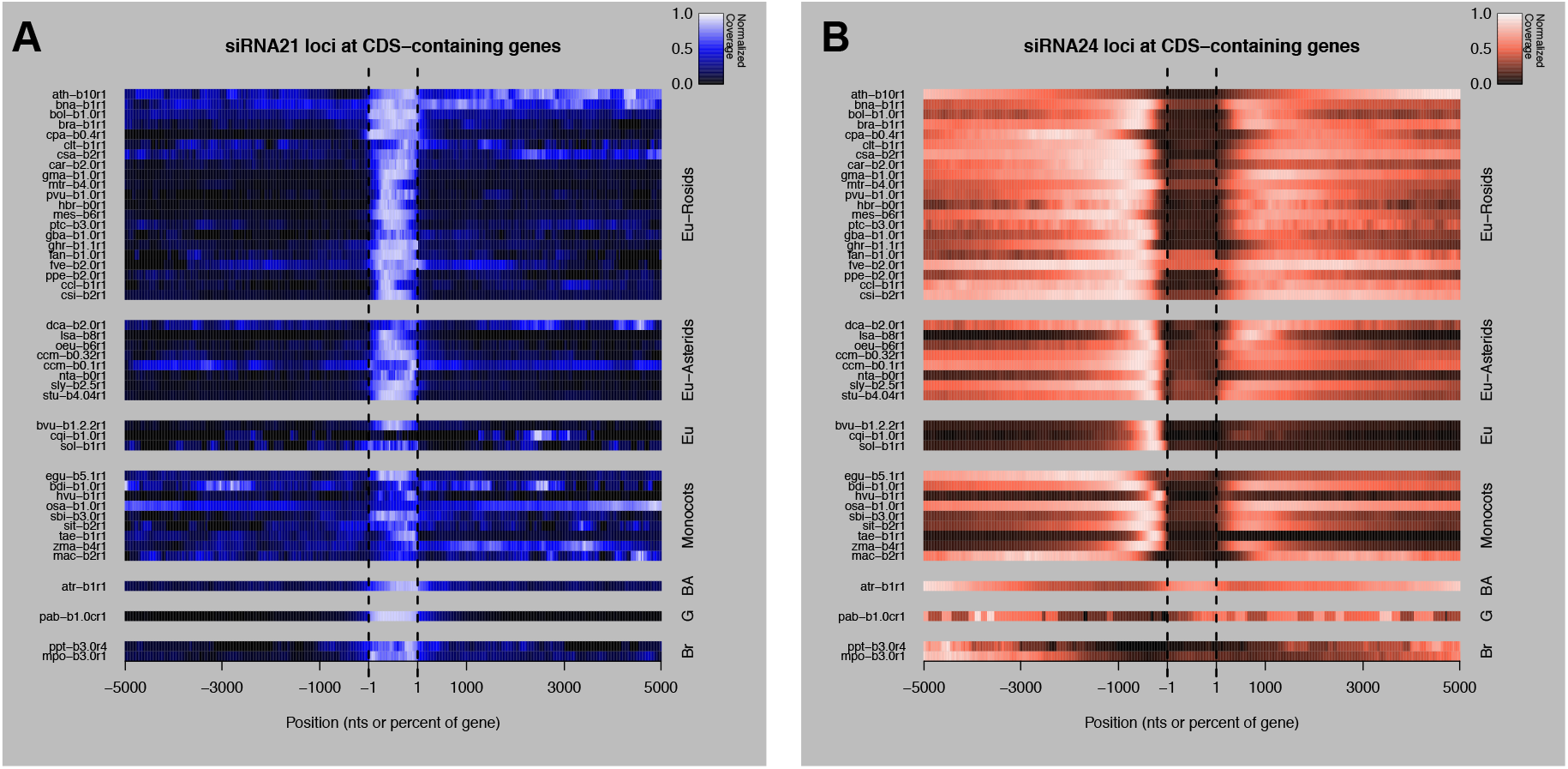
Associations of siRNA21 loci and siRNA24 loci with protein-coding genes. **(A)** Heatmap showing normalized coverage of protein-coding genes +/-5kb by siRNA21 loci. Each row is a given species (See Table 1 for species codes), grouped taxonomically. Eu: eudicots, BA: basal angiosperm, G: gymnosperm, Br: bryophyte. Negative and positive numbers are upstream and downstream regions, respectively (in nucleotides). The region from −1 to +1 represents the gene bodies, scaled to a uniform size of 1,000 nominal units of 0.1% each. **(B)** As in A, except for siRNA24 loci.

### Distribution of sRNAs in exons and introns of protein-coding genes

We then analyzed the distribution of sRNAs mapped to protein-coding genes, relative to the mRNA exons/introns and relative to the coding/non-coding strand of the mRNA (Figure 4). Although siRNA24 loci were generally depleted in mRNAs (Figure 3), their very large numbers still resulted in many overlaps, and therefore they were included in this analysis (Supplemental Fig. S6). For each species, we calculated the proportion of mRNAs that have 0% to 100% sRNAs mapped to the exons and the proportion of mRNAs that have 0% to 100% sRNAs mapped to the same strand of the mRNA. The proportions were plotted separately for mRNAs containing siRNA21 and siRNA24 loci. mRNAs containing siRNA21 loci showed a strong association with sRNAs arising from exons in the vast majority of the species (Figure 4A). These exonic 21 nucleotide siRNAs are most likely secondary siRNAs derived from the processing of the mRNAs. In contrast, in the mRNAs containing siRNA24 loci, sRNAs were primarily generated from introns in nearly all species (Figure 4B). Because 24 nucleotide siRNAs are known to be enriched in TEs, these intronic 24 nucleotide siRNAs could often be generated from intronic TE insertions. Some species showed a lesser association of siRNA24 loci with introns: this may be caused by differences in the annotation of TEs, which can sometimes be erroneously annotated as mRNAs. The siRNAs at both siRNA21 and siRNA24 loci typically originated from both strands of their associated genes (Figure 4C-D). This trend is consistent with processing from dsRNA precursors, as opposed to breakdown products from the mRNAs themselves.

**Figure 4.**
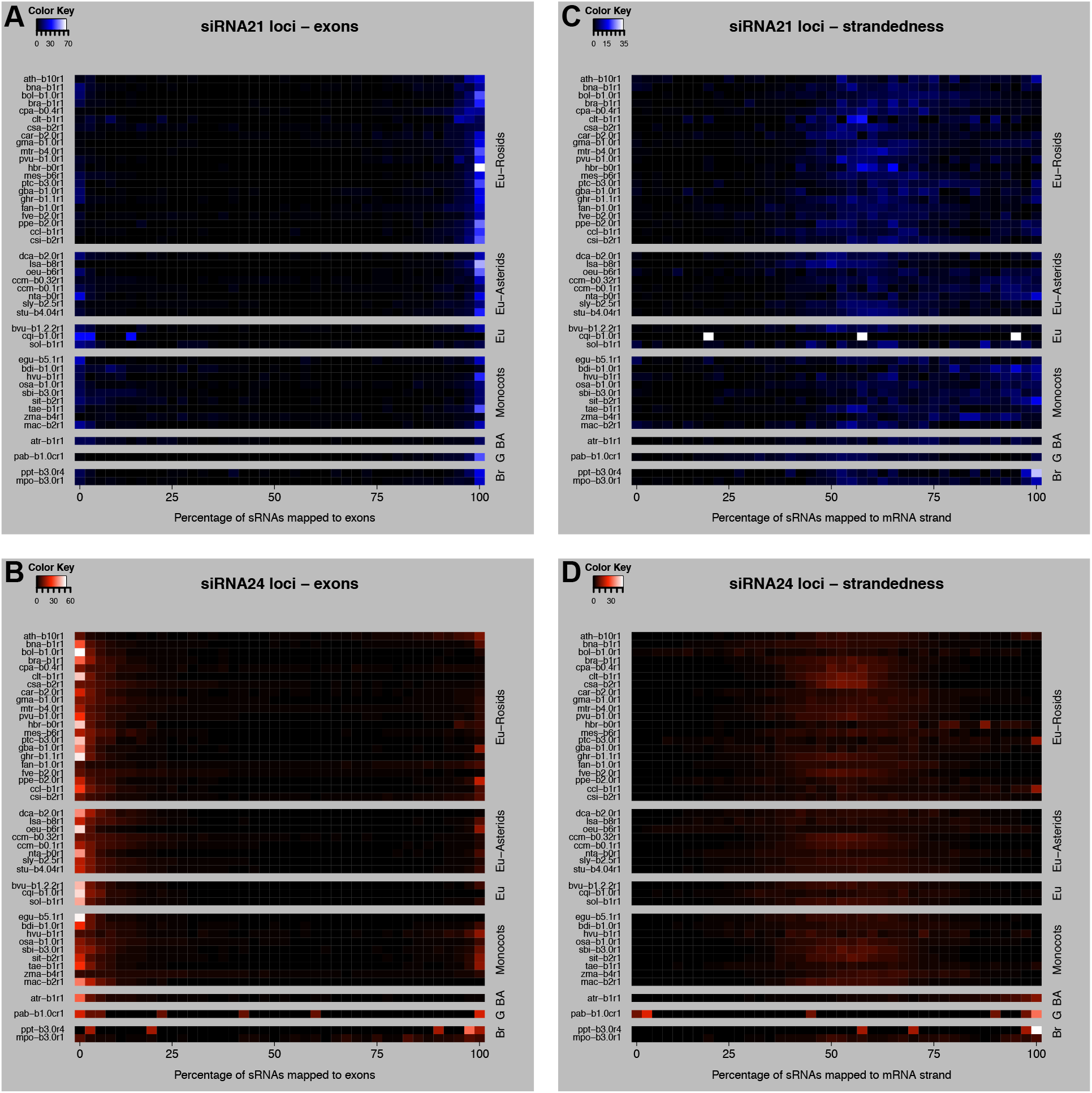
Distribution of sRNAs in the body region of protein-coding mRNAs. **(A)** Heatmap showing the proportion of mRNAs containing siRNA21 loci that have 0% to 100% of their aligned sRNAs mapped to their exons. 0% means all sRNAs map to introns, 100% means all sRNAs map to exons. **(B)** as in A, except for siRNA24 loci. **(C)** Heatmap showing the proportion of mRNAs containing siRNA21 loci with 0% to 100% of their aligned sRNAs mapped to the coding strand of the mRNA: 0% means all sRNAs map to the non-coding strand, 100% blue means all sRNAs map to the coding strand of the mRNA. **(D)** as in C, except for siRNA24 loci. Each row is a given species (See Table 1 for species codes), grouped taxonomically. Eu: eudicots, BA: basal angiosperm, G: gymnosperm, Br: bryophyte.

### Identity of genes associated with sRNA loci

To begin to understand the function of genes associated with sRNA loci, we performed GO enrichment analysis on the protein-coding genes that contained siRNA21 or siRNA24 loci, or siRNA24 loci in their 1 kb upstream region (Figure 5). For 38 of the 48 plant genomes, we were able to easily retrieve adequate GO annotations. These were used to perform Fisher’s Exact Test in Blast2GO in each species (FDR < 0.05). We plotted the frequency at which the GO terms were found enriched amongst the species to find conserved terms (Figure 5A). Enriched GO terms commonly found in at least ten species were considered to be well conserved, because at this number the frequency distribution inverted after gradually decreasing to zero. Genes containing siRNA21 and siRNA24 loci had respectively two and eight well conserved GO terms. In contrast, genes with siRNA24 loci within their 1 kb upstream region had no enriched GO term shared by ten or more species. The species distribution of the well conserved GO terms (Figure 5B) revealed that the “ADP binding” term was enriched in genes containing siRNA21 loci in rosids, asterids, in *A. trichopoda* and only in one monocot (wheat). Genes associated with the ADP binding function corresponded in all species with NB-LRR type disease resistance genes, which are known to produce secondary siRNAs in many species and only in barley and wheat amongst the monocots (Liu et al. 2014; Zhang et al. 2019). The “protein binding” term was also enriched in genes containing siRNA21 loci, but the genes associated with this term had heterogeneous and variable annotations between species, therefore no single common pathway was identified. Nevertheless, a few gene families in the “protein binding” group were commonly found amongst species, for example F-box genes, PPR-containing genes, kinases and SET domain containing genes. Genes containing siRNA24 loci had well conserved enriched GO terms mostly found in all clades and with different molecular functions (Figure 5B): “terpene synthase” and “heme binding” (mostly cytochromes P450 and other peroxidases) were the most conserved, followed by five others, including the “ADP binding” function. We hypothesize that the genes with these specific functions might be particularly frequent targets of intronic TE insertions silenced by 24 nucleotide siRNAs.

**Figure 5.**
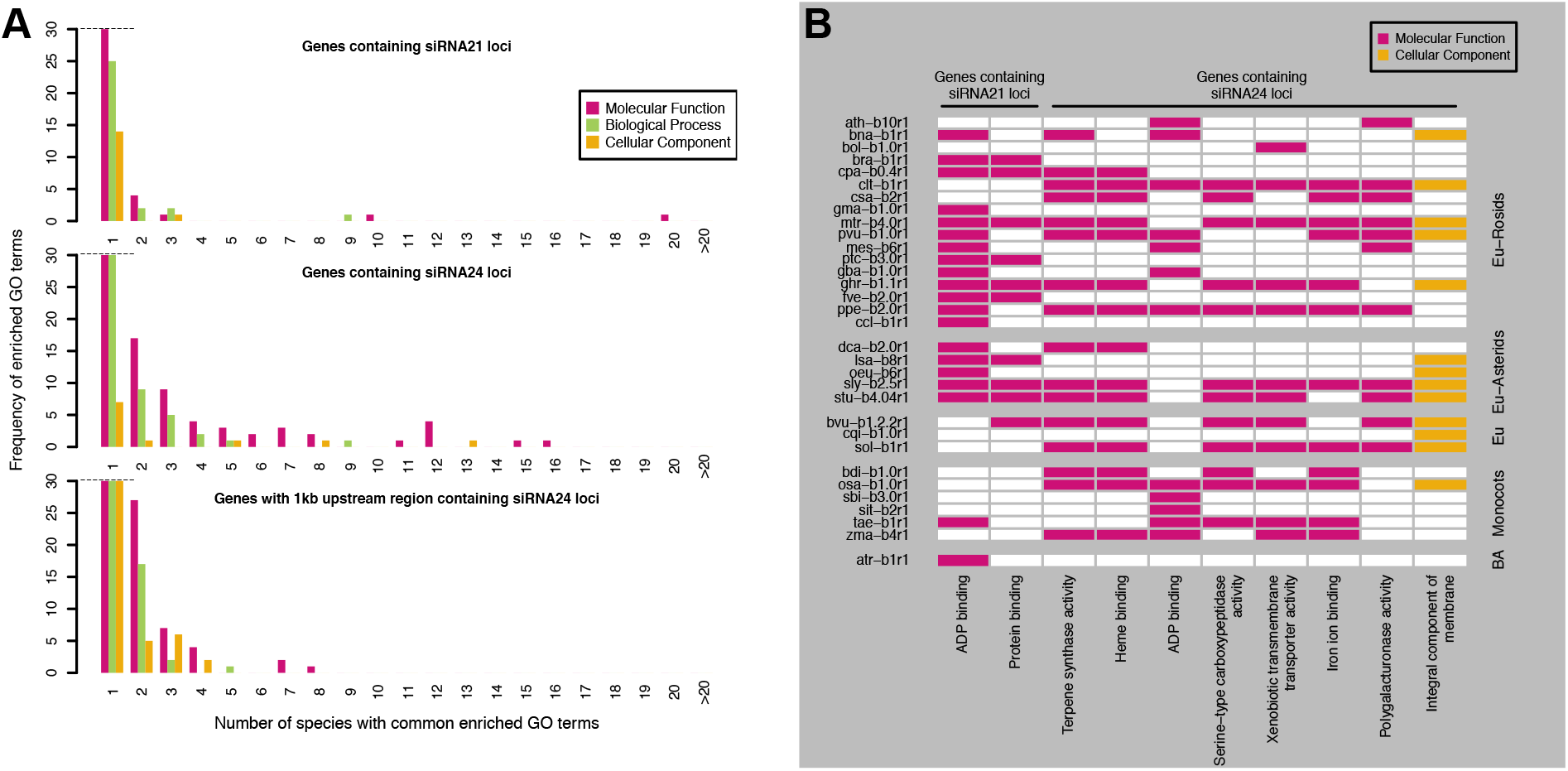
GO enrichment analysis of protein-coding genes associated with siRNA21 or siRNA24 loci. **(A)** Frequency of enriched GO terms in the 38 species analyzed (hbr-b0, tcc-b1.1, fan-b1.0, mdm-b3.0, csi-b2, ccm-b0.32, ccm-b0.1, can-b1.6, nta-b0 and hvu-b1 were excluded because no gene annotation or no GO annotation was available). **(B)** Species distribution of well conserved enriched GO terms, common to ten or more plant species (car-b2.0, egu-b5.1, mac-b2, pab-b1.0c, ppt-b3.0 and mpo-b3.0 were not displayed because they were not enriched in any of these terms). Each row is a given species (See Table 1 for species codes), grouped taxonomically. Eu: eudicots, BA: basal angiosperm.

### Disease resistance genes and other genes producing siRNAs in monocots

We further examined the nature of the genes containing exonic sRNA loci in monocots. This was of interest because the regulation of disease resistance genes by sRNAs in monocots has been described only in barley and wheat so far (Liu et al. 2014; Zhang et al. 2019). Genes containing exonic siRNA21 and siRNA22 loci were both studied, because within the monocots, maize produced high quantities of 22 nucleotide siRNAs (Figure 1), whose function is not well-understood. The genes were manually screened to discard those with stacks of sRNA reads mapped at only one or two unique positions, that could be alignment artifacts or miRNA-like sRNAs. Known miRNA matches, lowly expressed sRNA loci (< 1 RPM, reads per million), transposons and inverted repeats were also discarded. In total, 524 genes in the nine monocots were selected as containing robust siRNA21 and siRNA22 loci (Figure 6A, Supplemental Table S3). Maize was the only species where the majority of genic siRNA loci were siRNA22 loci; wheat also had some genic siRNA22 loci. This suggests that in maize and maybe wheat, the 22 nucleotide siRNAs could be a functionally active class of sRNAs in the regulation of genes, in addition to the 21 nucleotide siRNAs.

**Figure 6.**
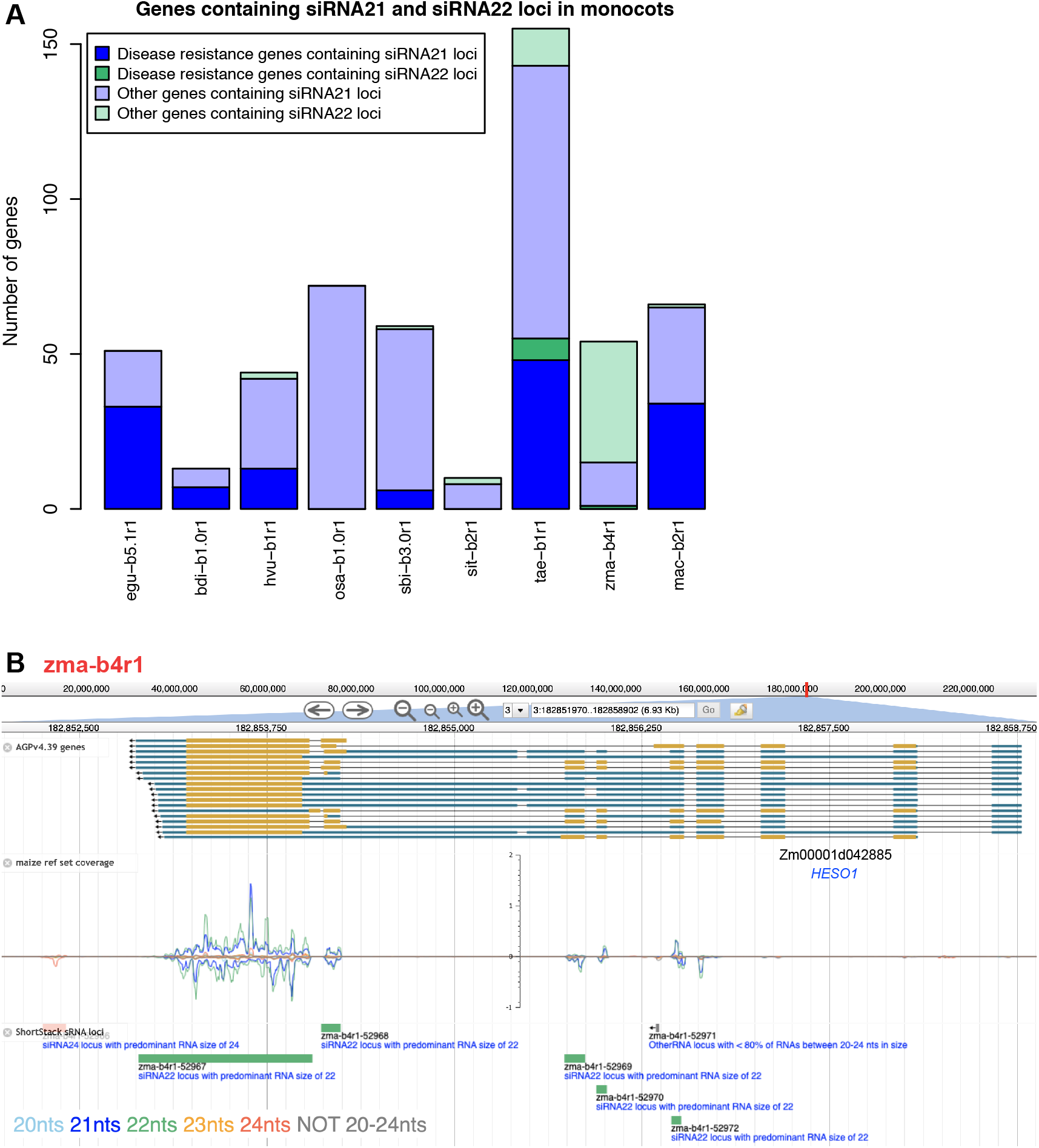
Genes containing siRNA21 and siRNA22 loci in nine monocot species. **(A)** Counts of genes containing siRNA21 and siRNA22 loci in monocots. **(B)** Screenshot of genome browser for maize *HESO1* (Zm00001d042885). Top row: mRNA structure: blue blocks for UTRs, yellow blocks for CDS and black lines for introns. Middle row: sRNA-seq coverage from the reference set across the gene. Bottom row: ShortStack sRNA loci annotation.

Evidence of sRNA expression from genes annotated as or having sequence homology with disease resistance genes, was found in seven species (Supplemental Table S3). Confirming previous reports, 13 disease resistance genes in barley and 48 in wheat contained siRNA21 loci. In oil palm and banana, 33 and 34 disease resistance genes, produced 21 nucleotide siRNAs, respectively. Disease resistance genes evolve rapidly by tandem duplications (Yang and Huang 2014), whose expression may be controlled by siRNAs. In the banana genome we found an example of this where two clusters of disease resistance genes, both on chromosome 3, contained 23 and 15 genes in tandem in a range of ∼137 and ∼130kb, respectively, that were sources of 21 nucleotide siRNAs. In *B. distachyon* and sorghum, we found seven and six resistance genes producing 21 nucleotide siRNAs, respectively, while in maize only one resistance gene produced 22 nucleotide siRNAs. In rice and foxtail millet there were no disease resistance genes associated with exonic 21 or 22 nucleotide siRNAs. This result suggests that the siRNA-mediated regulation of resistance genes could be conserved in a larger number of monocots than just barley and wheat but be selectively absent in some other monocots like rice.

Genes with different functions than resistance genes also contained siRNA21 and siRNA22 loci in monocots and a few were conserved in multiple species (Supplemental Table S3). Example of these genes include: *TAS3* genes, auxin responsive genes, kinase genes, genes encoding transport inhibitor response 1-like (TIR1-like) proteins, predicted E3 ubiquitin ligase genes, genes encoding or similar to DNA-directed RNA polymerases, two-component response regulators and methyl-CpG-binding domain-containing proteins. Genes participating in sRNA pathways were also found to be sources of siRNAs: *HEN1 SUPPRESSOR1* (*HESO1*, Figure 6B) and *AGO108* in maize, *DOMAINS REARRANGED METHYLTRANSFERASE 2* (*DRM2*) in rice, a predicted *AGO1B* in sorghum and three predicted copies of *AGO2* in wheat. As it is visible by the sRNA alignment coverage in *HESO1* (Figure 6B), sRNAs were expressed from multiple adjacent exons. This pattern of sRNA expression that reflects the mature mRNA structure was observed in many genes and strongly suggests that these exonic 21 and 22 nucleotides sRNAs are secondary siRNAs, originated from the processing of the mRNA by a DCL protein.

### Analysis of sRNA conservation across plant species

Annotated sRNA loci were grouped into putative families based on the sequences of the most abundant single sRNA (the ‘major RNA’) produced by each locus (Supplemental Table S4). Loci were considered to be members of the same family if the sequences of their major RNAs had up to two mismatches with each other; these criteria are similar to those commonly used to group *MIRNA* loci into families. Most of the resulting families (1,556,834; 85.3%) had only a single locus (Figure 7A) and relatively few families (38,794; 2.1%) were present in more than a single species (Figure 7B). Even fewer families (1,968; 0.1%) were present in more than one major taxonomic group (Figure 7C). In general, the proportions of MIRNA, nearMIRNA, and siRNA21 loci were higher for more extensively conserved families (Figure 7D-F); at the most extreme levels of conservation, MIRNA loci and siRNA21 loci predominated.

**Figure 7.**
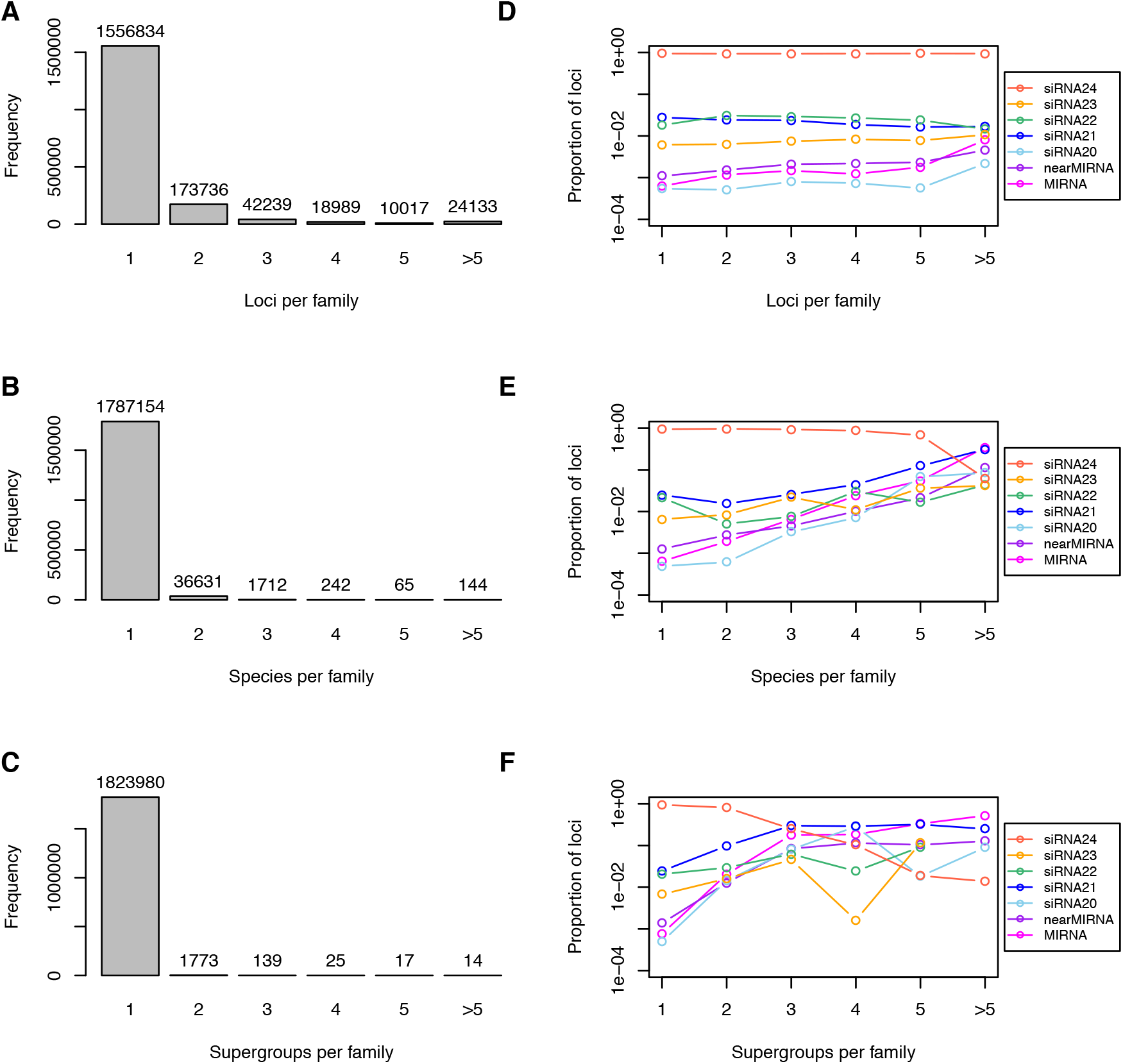
Conservation of sRNA loci in plants. **(A)** Frequency distribution of number of sRNA loci per putative sRNA family. **(B)** Frequency distribution of number of distinct plant species per putative sRNA family. **(C)** Frequency distribution of number of plant ‘supergroups’ per putative sRNA family. Supergroups defined in this study are: rosids, asterids, other eudicots, monocots, basal angiosperms, gymnosperms, and bryophytes. **(D)** Proportions of types by number of loci per putative sRNA family. **(E)** Proportions of types by number of distinct plant species per putative sRNA family. **(F)** Proportions of types by number of plant ‘supergroups’ per putative sRNA family.

## Discussion

### A public resource on sRNAs for the scientific community

We created an extensive resource for a large number of plant genomes that allows users to freely and easily retrieve, visualize and analyze sRNA loci, including not only miRNA annotations but also siRNA annotations. Our research extended into non-model systems, including many species of horticultural importance. For three economically important plants, spinach, carrot and cacao, we annotated for the first time miRNA and siRNA loci. Recently, a study published the first miRNA annotation in carrot using high throughput sequencing, but the siRNAs were not examined (Bhan et al. 2019). In many published works, sRNA-seq is used to annotate and profile miRNAs but not individual siRNA loci. We used the vast amount of available sRNA-seq datasets to exploit all this unrevealed information and annotate the entire population of sRNAs in 48 plant genomes.

Our database and analyses are limited by the quality and quantity of the available genomic annotations and sRNA-seq data. For example, sRNAs expressed in specific tissues/cell types or growth conditions that were not represented in our sRNA-seq dataset are by consequence absent from our reference annotations. This might be the case for reproductive phasiRNAs (Zhai et al. 2015; Fei et al. 2016; Xia et al. 2019): for most of the analyzed species we did not have any or enough sRNA-seq libraries from male reproductive tissues to allow the specific annotation of this sRNA population. For this reason, we did not investigate the reproductive sRNAs in our work. Our database has the potential to be expanded in the future to include new plant genomes, new annotations and new sRNA-seq data that are of interest for the plant biology community.

Overall, we created a resource that will be useful for future sRNA studies. Thanks to the standard annotation and classification methods followed for all genomes, our sRNA annotations and alignments can be directly visualized or downloaded from our web-server and compared between species. Our web-server is a practical way to quickly interrogate existing plant sRNA data in a usable format and will enable scientists to rapidly search for evidence of sRNA expression in specific regions in a species or investigate the conservation of single sRNA sequences across species.

### Multiple protein-coding gene families are sources 21 nucleotide siRNAs in dicots and monocots

The best characterized case of protein-coding genes generating secondary siRNAs are the disease resistance genes, whose expression is kept under control by secondary siRNA production to avoid fitness loss (Yang and Huang 2014). We confirmed expression of 21 nucleotide siRNAs from exons of resistance genes in the rosid and asterid clades and expanded the number of monocot species that also showed this evidence, suggesting that this pathway might be more broadly conserved than what is known. In none of the three studied caryophyllales species the protein-coding genes containing siRNA21 loci were enriched in the GO:ADP binding term, characteristic of resistance genes. This could result from incomplete gene/GO annotations in spinach, sugar beet and quinoa, missing real resistance genes. Alternatively, these secondary siRNAs might be reduced in caryophyllales because the number of disease resistance genes in this clade is lower compared to the typical expansion of this gene family in rosids and asterids or because in caryophyllales, specific subfamilies of resistance genes have expanded that might be differentially regulated (Dohm et al. 2014; Xu et al. 2017; Funk et al. 2018).

In the literature, the number of known protein-coding genes producing secondary siRNAs in monocots is smaller than in dicots. Accordingly, from our analyses, the enrichment of siRNA21 loci in protein-coding genes was less evident in monocots compared to dicots and also the tendency of 21 nucleotide siRNAs to map to exons was smaller in monocots. For these reasons, we decided to manually screen the monocot species for evidence of 21 nucleotide siRNA production from protein-coding genes. We described a number of gene families, more or less conserved in the nine monocots, that produced 21 nucleotide siRNAs, and also 22 nucleotide siRNAs in maize and wheat. In many cases, the siRNAs were expressed specifically from multiple adjacent exons, supporting the hypothesis that they are secondary siRNAs processed from mature mRNAs. Some of the genes found were previously described as sources of secondary siRNAs in other species, for example kinase genes (Zheng et al. 2015; Reyes-Chin-Wo et al. 2017), TIR1-like genes (Si-Ammour et al. 2011; Seo et al. 2018; Xia et al. 2015a) and *AGO2* (Arikit et al. 2014). In addition to *AGO2*, there are more genes participating in siRNA biogenesis and function that are themselves known targets of siRNA regulation: *DCL1* (Xie et al. 2003; Hu et al. 2015b; Xia et al. 2014), *DCL2* (Zhai et al. 2011; Arikit et al. 2014), *AGO1* (Vaucheret et al. 2006) and *SUPPRESSOR OF GENE SILENCING 3* (Arikit et al. 2014). We found evidence of siRNA expression from four additional genes involved in siRNA pathways: in maize, from *AGO108* (also named *AGO5d*), highly expressed in ears but not well functionally characterized (Zhai et al. 2014), and *HESO1*, a nucleotidyl transferase that uridylates unmethylated sRNAs to trigger their degradation (Zhao et al. 2012); in sorghum, from a predicted *AGO1B* and in rice from *DRM2*. *DRM2* is a known target of miR820 in rice (Nosaka et al. 2012), which could be the trigger miRNA for the production of the observed 21 nucleotides siRNAs. We reported many more genes in monocots that spawned 21 or 22 nucleotide long siRNAs, belonging to different families. These genes represent an interesting set to research in the future to better characterize the nature of genic siRNAs. The next obvious step will be searching for possible miRNA triggers and examining the phasing pattern of siRNA expression in each specific gene, to confirm that these siRNAs are secondary siRNAs.

### Different hypotheses on 22 nucleotide siRNA functions

We found that asterids consistently had considerable proportions of siRNA22 loci, while in the other clades, only certain species (soybean, cassava and maize) had this same trend. There are several hypotheses that could explain the presence of 22 nucleotide siRNAs in a genome: they could originate from *MIRNA* or *MIRNA*-like loci that were missed by our annotation method, from endogenous direct or inverted repeats (Kasschau et al. 2007), or from protein-coding genes, as we observed in maize. Alternatively, these siRNA22 loci could express siRNAs involved in the non-canonical RdDM pathway to silence active TEs (Matzke and Mosher 2014), as it was proposed for maize (Nobuta et al. 2008). Active retrotransposons have been described in asterids, for example the Tto1 element or the Tnt1 element, which has many copies that are still transcriptionally active in tobacco (Casacuberta et al. 1997) and lettuce (Mazier et al. 2007). In this hypothesis, what still remains unclear is why we observed expression of 22 nucleotide siRNA most often in the asterids and not in the grasses, where retrotransposon transcription is very prevalent (Vicient et al. 2001). If the 22 nucleotide siRNAs come from active retrotransposons, then the ability to detect their expression could depend on the specific samples analyzed, because retrotransposons are only active during certain stages of plant development or stress conditions (Flavell et al. 1992). Lastly, 22 nucleotide siRNAs could target Endogenous Viral Elements, virus segments that are integrated in the host genome, that form inverted repeats (Pooggin 2018). To understand the role of the siRNA22 loci, the next step in future research will be the genome-wide profiling of the genomic regions where these loci map, discriminating between genes, intergenic regions and different classes of TEs.

### Roles of 24 nucleotide siRNAs in regulating protein-coding gene expression

We assumed that the distribution of the total sRNA loci across the chromosome length reflected the distribution of the siRNA24 loci, because these accounted for the vast majority of loci in angiosperms. *A. thaliana* and Chinese cabbage are two of the few species where siRNA24 loci and gene densities were negatively correlated. In both species, siRNA regulation of TEs near genes was previously linked to lower expression of the genes (Hollister et al. 2011; Woodhouse et al. 2014). It would be informative to test if the same link occurs in the other species with inverse correlation between siRNA24 locus and gene densities, like sweet orange. Differences in siRNA24 locus distribution and influence on gene expression might be directly explained by differences in TE composition between genomes. Accordingly, it was previously suggested that the transcription of gene networks can be balanced by the genome distribution of TEs (Freeling et al. 2015), which can be highly variable among species (Vicient and Casacuberta 2017). In many cases, a few TE families have increased their copy number in one lineage (Baidouri and Panaud 2013). For example, a single type of LTR retrotransposon is responsible for most of the hot pepper genome expansion (Park et al. 2012).

The angiosperms analyzed were strongly enriched in siRNA24 loci in the 5’-proximal regions upstream of protein-coding genes. In *A. thaliana*, this distribution was much less strong but the enrichment of siRNA24 loci in the 5’ upstream region compared to the gene body region was still evident. The function of siRNA24 loci at these sites has been widely studied in maize: near genes, 24 nucleotide siRNAs engage RdDM, blocking the spread of open, active chromatin into adjacent transposons (Li et al. 2015). In addition to silencing TEs, the RdDM activity near genes in *A. thaliana* can also affect the expression levels of the genes (Zheng et al. 2013; Zhong et al. 2012), likely by changing the chromatin landscape at gene promoters and influencing the ability of transcription factors to bind to the promoters and stimulate transcription. In maize on the contrary, no obvious direct effects on gene expression were detected as a consequence of the loss of gene proximal 24 nucleotide siRNAs (Lunardon et al. 2016). Finally, in *A. thaliana*, it has been speculated that the RdDM activity near genes can influence their expression by inhibiting interactions between the promoters and their potential distant regulatory elements (Rowley et al. 2017). Similarly, most angiosperms were also enriched in siRNA24 loci at the 3’-proximal regions downstream of genes, where the RdDM activity seems to reduce the readthrough transcription by Pol II into neighboring genes or TEs (Erhard et al. 2015).

When siRNA24 loci were found inside protein-coding genes, they were mostly in introns. A few gene families were most commonly targeted by 24 nucleotide siRNAs in both dicots and monocots. Two possible reasons might explain why these specific genes were a common target of 24 nucleotide siRNAs. On one side, families like disease resistance genes evolve rapidly, creating high numbers of partial genes and pseudogenes (Luo et al. 2012) that might be suppressed by the activity of 24 nucleotide siRNAs (Kasschau et al. 2007). This could also be the case of polygalacturonases that are encoded by a large gene family. An accurate study of the protein-coding gene annotations, precisely separating genes from pseudogenes, would be necessary to verify this hypothesis. On the other side, gene families like disease resistance genes control adaptive responses to the environment, making them frequent targets of TE transposition events (Quadrana et al. 2016). Although the majority of TE insertions in genes are deleterious, they can be advantageous and therefore be retained as source of variability, which is essential in environmental response genes to adapt to the ever-changing environment. As a consequence, new TE insertions are overrepresented in genes that respond to environmental stresses (Grover et al. 2003; Miyao et al. 2003). Also in cytochrome P450s, a family known to participate in stress responses, frequent TE insertions were described as a strategy for variability (Chen and Li 2007) and this could explain why these genes were frequent targets of 24 nucleotide siRNAs. Likewise, serine-type carboxypeptidases, which participate in protein degradation, and xenobiotic transmembrane transporters, which work in xenobiotic detoxification pathways together with cytochromes P450, both play pivotal roles in plant defense responses and therefore could be frequent targets of TE insertions controlled by 24 nucleotide siRNAs. To verify if the intronic 24 nucleotide siRNAs influence the regulation of the genes that they target, it will be informative in the future to examine mutants lacking the production of 24 nucleotide siRNAs and observe if these gene families tend to be altered in their expression.

### Conservation of siRNAs

The sequence comparison of the most abundant sRNA expressed from each locus revealed a very low level of conservation of siRNAs across species, not just between distant species but also between close relatives. Studying the conservation of siRNAs is complicated by the fact that the siRNA population can vary substantially between different organs of the same plant species (Ha et al. 2009). Nonetheless, our result is in line with previous observations (Ma et al. 2010). If we consider plants that all have a strong peak of 24 nucleotide siRNAs and have a functional RdDM pathway, the genomic TE composition and organization can significantly differ between different species and even between different varieties of the same species (Brunner et al. 2005; Quadrana et al. 2016). This might explain why the individual siRNA sequences that target the TEs are also poorly conserved. Much of our knowledge regarding sRNAs comes from model plants like *A. thaliana*, which has a low amount of TEs that are not active in wild-type plants. Crop genomes, instead, have high TE loads and some TEs are active in wild-type genetic backgrounds in maize and rice (Jiang et al. 2003; Nakazaki et al. 2003; Lisch 2012). Due to these differences it is important to study sRNAs in non-model systems, because lineage-or species-specific sRNAs might be associated to traits that other plants lack or have not evolved (Chen et al. 2018).

## Methods

### Plant material and sRNA sequencing

Leaves of *Theobroma cacao* (line Scavina 6) were kindly provided by Dr. M. Guiltinan of The Pennsylvania State University, from plants grown in greenhouse conditions. The tips of leaves at the immature green leaf stage were collected. *Daucus carota* (cultivar ‘Burpee’) was grown in a growth room at 22°C, 16h light 8h dark regime and leaves and roots from 5- and 6-week-old plants, respectively, were sampled. *Spinacia oleracea* Sp75 inbred line seeds were kindly provided by Dr. Z. Fei of the Boyce Thompson Institute, Cornell University, and grown in a growth room at 22°C, 16h light 8h dark regime. Leaves from 3- and 5-week-old plants were collected. *Zea mays* B73 inbred line seeds were germinated on ProMix B, then transferred to soil in pots and grown in greenhouse conditions with occasional Osmocote fertilization. The fifth and the sixth leaves from V5 plants, mature pollen and 21-27 DAP (days after pollination) embryo tissue were collected from a pool of plants. All samples were flash frozen in liquid nitrogen, stored at −80°C and then ground with liquid nitrogen cooled mortar and pestle. For carrot, spinach and maize, the RNA was extracted with Tri-reagent (Sigma) as per manufacturer instructions, adding a second sodium-acetate–ethanol precipitation and ethanol wash step. For cacao, the RNA was extracted with PureLink Plant RNA Reagent (Life technologies) following manufacturer’s suggestions. Sequencing libraries were prepared using the NEB Next sRNA-seq library preparation kit for Illumina (NEB, E7300S) following manufacturer’s suggestions. Reactions were purified and size selected for sRNAs 15-40nt in length by PAGE. Extracted bands were quantified by qPCR and quality-controlled by high-sensitivity DNA chip (Agilent). Sequencing was performed on a HiSeq2500 (Illumina) in rapid run mode (50 nucleotides, single-end, single barcode) by the Penn State genomics core.

### sRNA-seq data processing

sRNA-seq raw fastq files were downloaded from SRA and GEO databases (Supplemental Table S1). The libraries were processed to remove the 3’ adapter with cutadapt (Martin 2011) (cutadapt -a *3’_adapter_sequence* --discard-untrimmed -m 15 -o *output_file.fastq input_file.fastq*). Reads containing the 5’ adapter were removed with cutadapt (cutadapt -g *5’_adapter_sequence* --discard-trimmed -m 15 -o *output_file.fastq input_file.fastq*). Low quality reads were discarded with FASTX-Toolkit (Gordon and Hannon 2010) (fastq_quality_filter -q 20 -p 85 -Q 33 -v -i *input_file.fastq* -o *output_file.fastq*). Finally, reads quality was checked with FastQC (Andrews 2010): if additional sequencing adapters were overrepresented amongst reads, they were eliminated from the fastq files with a custom Perl script.

### Pipeline to create reference sRNA loci annotations

For each species, the reference annotation of sRNA loci was created with the following steps. Each individual library was aligned to the genome (see https://plantsmallrnagenes.science.psu.edu for list of genome assemblies used) using ShortStack v3.8.1 (Axtell 2013b; Johnson et al. 2016) with default parameters. Libraries with less than 2 million mapped reads were discarded. Clusters of sRNAs were *de novo* identified in each library independently with ShortStack (ShortStack --bamfile *alignment_file.bam* --mincov 2rpm --genomefile *genome_file.fa*). The sRNA clusters files from all libraries of the same species were intersected with the bedtools function ‘multiIntersectBed’ (Quinlan and Hall 2010) with default parameters. Only genomic intervals with annotated sRNA clusters common to at least three libraries were kept and merged with bedtools, with 25 nucleotides as maximum distance allowed between the intervals to be merged into sRNA loci (mergeBed -d 25 -i *input_intervals_file.bed* > *output_merged_intervals_file.bed*). sRNA loci with length < 15 nucleotides were removed with a custom Perl script. Finally, sRNA loci whose expression was < 0.5 RPM in all libraries were also removed. The sRNA loci that were selected after applying these filters represented the reference annotation for each species.

### Analysis of sRNA loci occupancy relative to protein-coding genes

Locations of protein-coding genes were determined from public GFF3 files from each genome. Intergenic regions were calculated using bedtools ‘complement’, computationally cut in half, and associated with their nearest protein-coding genes using bedtools ‘closest’. The regions were marked as upstream or downstream based on the orientation of their nearest flanking gene. Per-nucleotide overlap between upstream, downstream, and gene-body regions vs. small RNA loci were calculated using bedtools ‘overlap’. The lengths of gene-bodies were scaled to 1,000 arbitrary units (each such unit is 0.1% of the gene length). Coverage was summarized in 25 nucleotides / unit bins, and normalized to a scale of 0 to 1, where 1 represented the maximum fraction occupancy observed in that genome.

### Analysis of sRNA distribution in exons and introns of protein-coding mRNAs

Only protein-coding mRNAs having at least one intron and overlapping with siRNA21 and siRNA24 loci were studied. Each mRNA was either classified as containing siRNA21 or siRNA24 loci: in case of overlap with both siRNA21 and siRNA24 loci, the longest sRNA locus was considered. The number of sRNAs mapped to exons and to the same strand of protein-coding mRNAs containing one or more introns were calculated with the bedtools function ‘coverageBed -counts’ (parameters added for exons: ‘-F 1’; for the same strand: ‘-F 1 -s’). The number of sRNAs mapped to introns and to the opposite strand of the mRNAs were also calculated for the final ratios (parameters added for the opposite strand: ‘-F 1 -S’). The percentage of sRNAs mapped to exons was calculated based on the ratio ‘number of reads mapped to exons / (number of reads mapped to exons + number of reads mapped to introns)’. The percentage of sRNAs mapped to the same strand of the mRNA was calculated based on the ratio ‘number of reads mapped to the same strand / (number of reads mapped to the same strand + number of reads mapped to the opposite strand)’. Here and in the other analyses of siRNAs, sRNA loci classified as MIRNA and nearMIRNA or whose most abundant sequence had a perfect match with a high-confidence plant miRNA hairpin annotated in miRBase v22 (Kozomara and Griffiths-Jones 2014) were not included.

### GO enrichment analysis

Protein-coding genes were classified as containing siRNA21 or siRNA24 loci and as flanked in their 1kb upstream region by siRNA21 or siRNA24 loci: when the same gene/upstream region overlapped with both siRNA21 and siRNA24 loci, the longest sRNA locus determined the classification. The GO enrichment analysis was performed with Blast2GO (Götz et al. 2008), using the Fisher’s Exact Test with default parameters (FDR < 0.05). Only the species for which we were able to retrieve a GO annotation were analyzed, this excluded: hbr-b0, tcc-b1.1, fan-b1.0, mdm-b3.0, csi-b2, ccm-b0.32, ccm-b0.1, can-b1.6, nta-b0 and hvu-b1.

### Analysis of genes containing siRNA21 and siRNA22 loci in monocots

To find all genes containing siRNA21 and siRNA22 loci in exons we used bedtools (intersectBed -wao -F 0.75 -a *exons_file.gff3* -b *sRNA_loci_file.gff3* > *output_intersection_file.txt*). When the same gene contained both siRNA21 and siRNA22 loci, if it contained a greater number of siRNA21 loci than siRNA22 loci it was classified as containing siRNA21 loci. In case there were the same number of siRNA21 and siRNA22 loci, the gene was classified based on the longest locus. The description of the genes (Supplemental Table S3) was copied from the gene annotation files retrieved from the same online resources used for the genome sequences (see https://plantsmallrnagenes.science.psu.edu for sources of genomes and gene annotations files). For species without available gene annotations, the function of the genes was predicted using BLAST (Camacho et al. 2009) on the gene sequence and considering the best result.

### Data access

All sRNA-seq libraries used, published and newly generated, are available in GEO and SRA; see Supplemental Table S1 for accession numbers. All data and analyses are hosted at https://plantsmallrnagenes.science.psu.edu.

## Acknowledgements

We thank the Penn State Genomics Core Facility for small RNA sequencing services, and the Eberly College of Science IT office for providing server hosting services for this project. We thank Matthew Jones-Rhoades for supervision and mentoring of undergraduate researchers and for insightful comments on the manuscript. This work was supported by an award from the US National Science Foundation (Award 1339207) to MJA.

## Author Contributions

AL and MJA generated most primary annotations and most analyses, with some primary annotations also contributed by SP. The project website was developed by MJA. NRJ, EH, TP, and CC contributed small RNA sequencing results. The manuscript was written by AL and MJA with input from NRJ and CC.

## Supplemental Tables

Supplemental Table S1. sRNA-seq libraries that were used as components in reference sets. Format: comma-separated values (csv).

Supplemental Table S2. Small RNA-producing loci from 48 plant genomes. Gzip-compressed, tab-separated text file. Note that the project website (http://plantsmallrnagenes.science.psu.edu) has the same data with search functions, more details, and alternative formats.

Supplemental Table S3. List of genes containing siRNA21 and siRNA22 loci in the nine monocot species studied.

Supplemental Table S4. Grouping of small RNA loci into putative families based on sequences of most abundant sRNA sequence. Gzip-compressed, tab-delimited text file.

## Supplemental Figures

**Supplemental Fig. S1.**
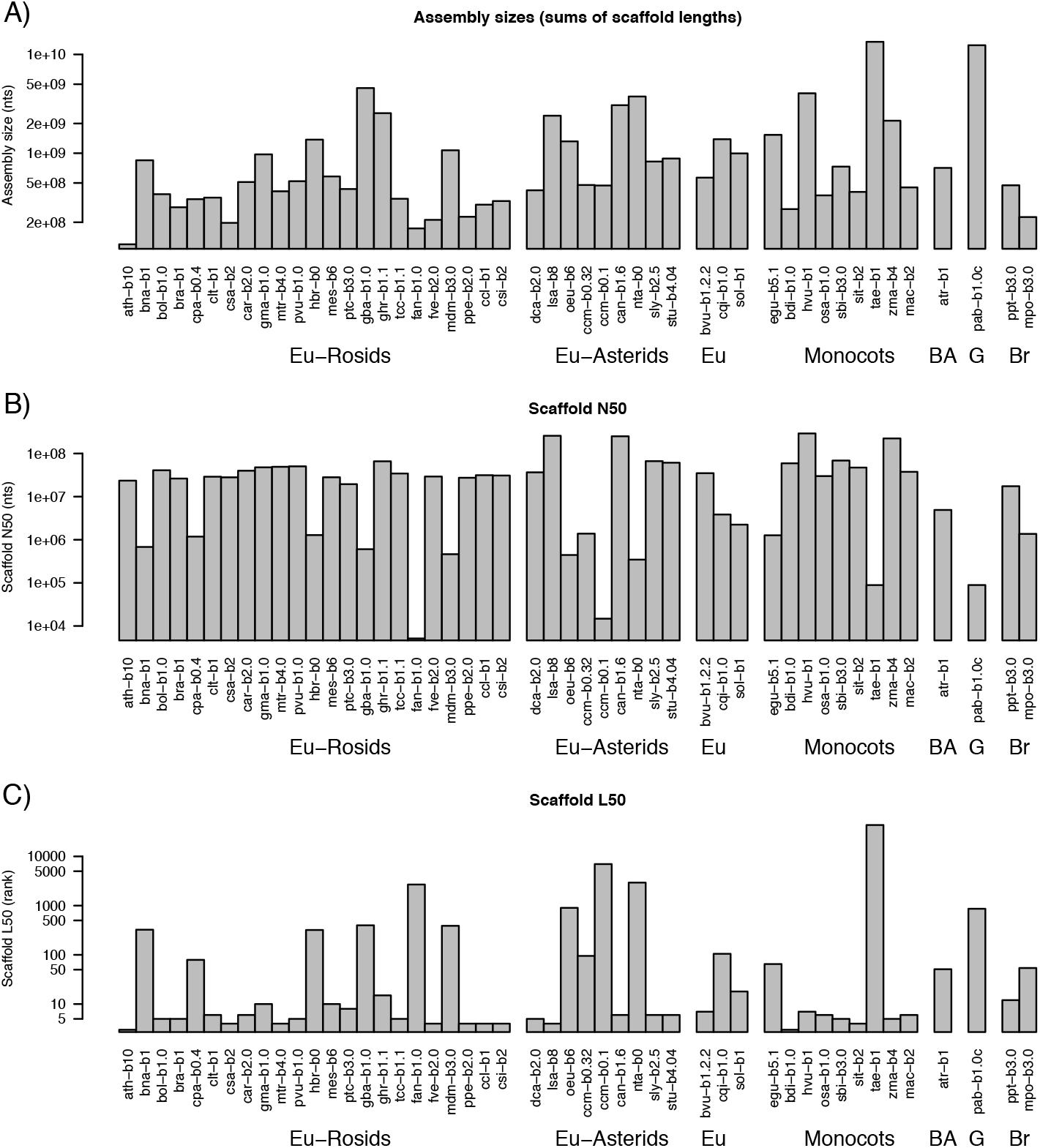
Properties of genome assemblies used in this study. See Table 1 for species codes. Eu: eudicots, BA: basal angiosperm, G: gymnosperm, Br: bryophyte. **(A)** Genome assembly sizes (log_10_ scale, nucleotides). **(B)** Scaffold N50 lengths (log_10_ scale, nucleotides). **(C)** Scaffold L50 ranks (log_10_ scale).

**Supplemental Fig. S2.**
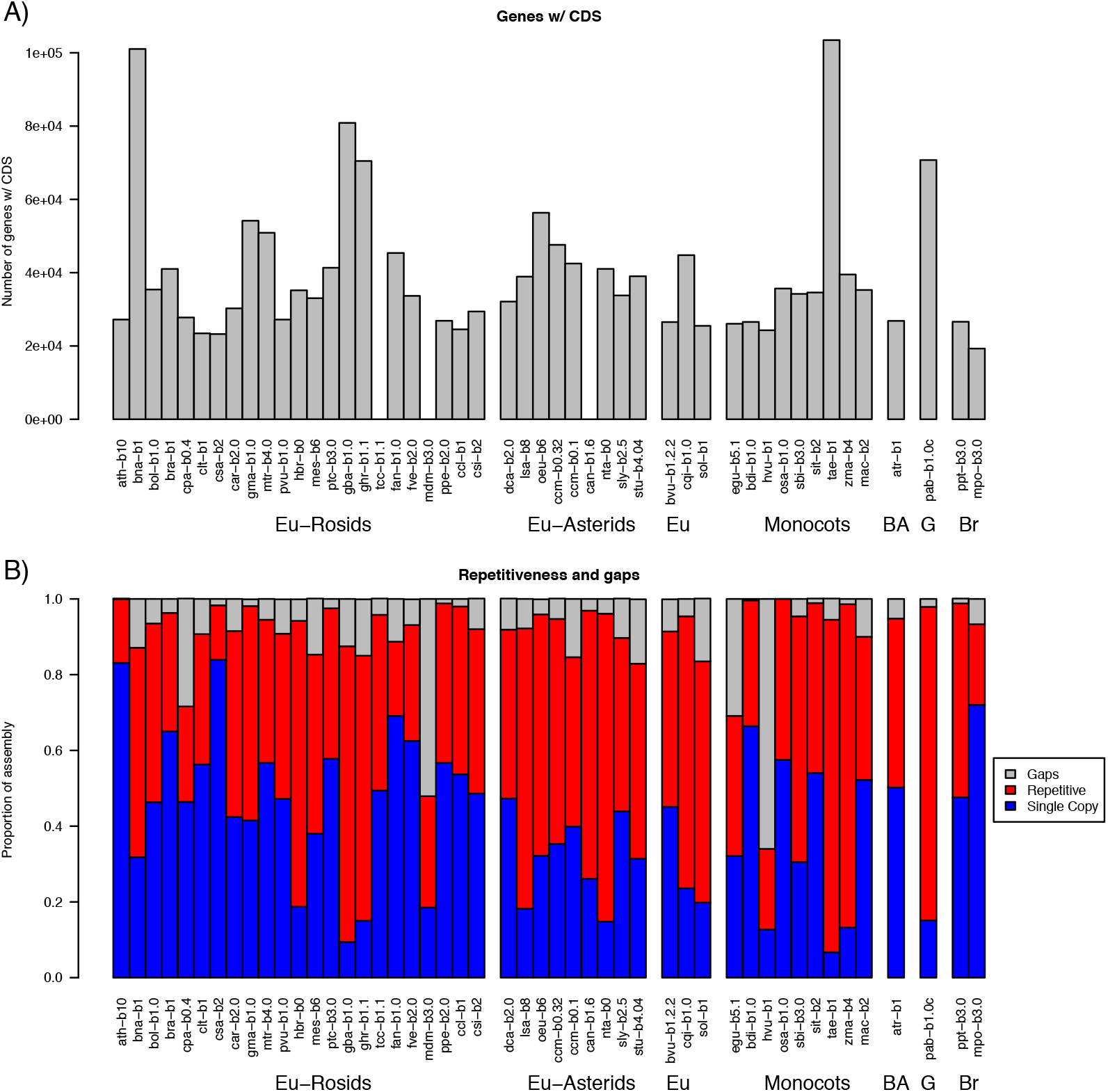
Properties of gene annotations and genome assemblies used in this study. See Table 1 for species codes. Eu: eudicots, BA: basal angiosperm, G: gymnosperm, Br: bryophyte. **(A)** Counts of annotated genes that contain one or more CDS features. Note: data were unavailable for three assemblies (tcc-b1, mdm-b3.0, can-b1.6). **(B)** Repetitiveness, gaps, and single-copy regions, defined by k-mer analysis of genome assemblies with k=24. K-mers containing one or more ambiguous character were tallied as gaps. Non-ambiguous k-mers present more than once in the assembly were tallied as repetitive.

**Supplemental Fig. S3.**
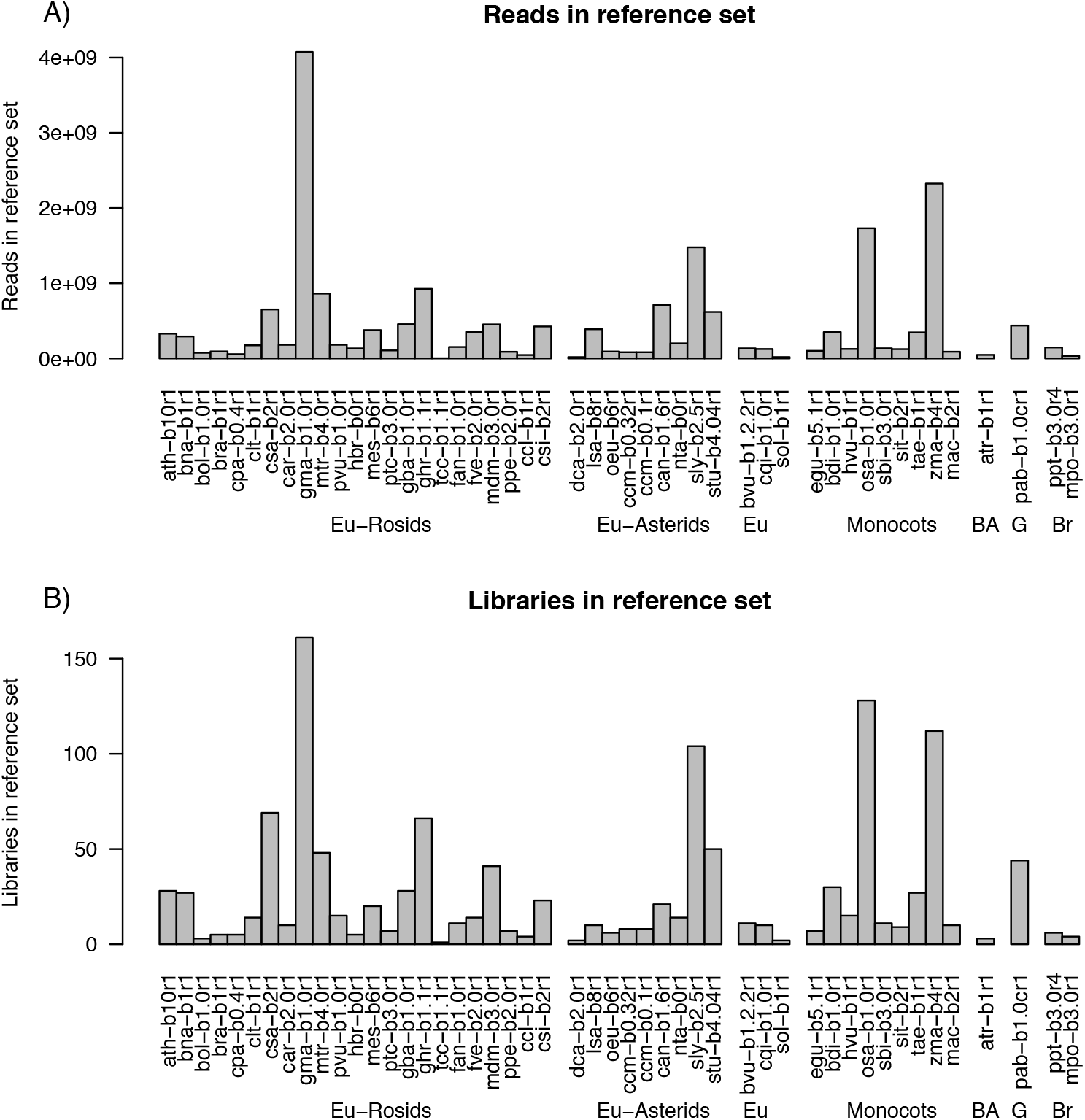
Summary of sRNA-seq libraries and reference sets. See Table 1 for species codes. Eu: eudicots, BA: basal angiosperm, G: gymnosperm, Br: bryophyte. **(A)** Total number of sRNA-seq reads in reference sets. **(B)** Number of individual sRNA-libraries in reference sets.

**Supplemental Fig. S4.**
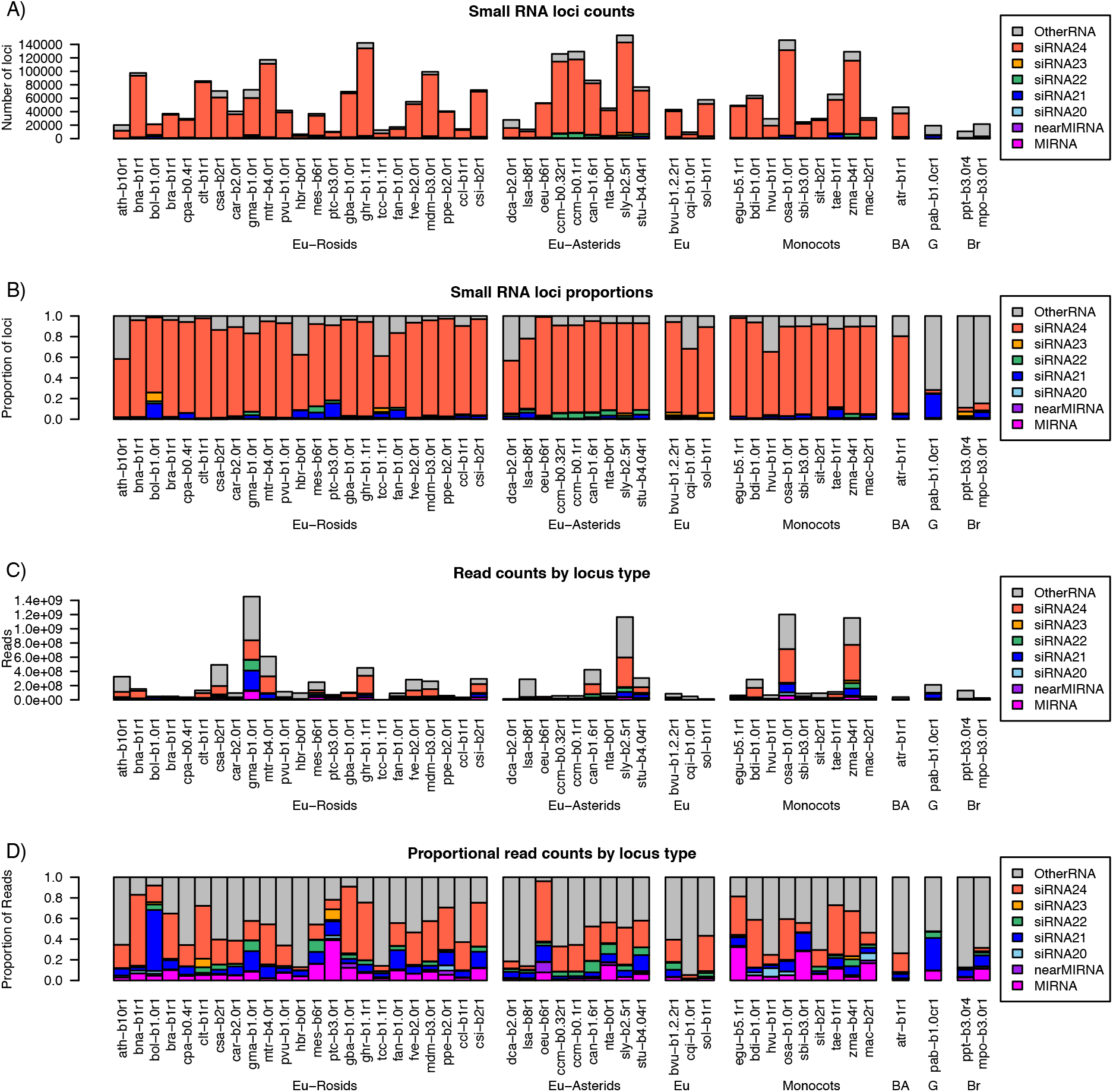
Summary of annotated sRNA loci, by species and locus type, including the category ‘OtherRNA’. See Table 1 for species codes. Eu: eudicots, BA: basal angiosperm, G: gymnosperm, Br: bryophyte. **(A)** Counts of annotated loci. **(B)** Proportions of annotated loci. **(C)** Total counts of aligned small RNAs in reference sets. **(D)** Proportions of small RNA total read counts in reference sets.

**Supplemental Fig. S5.**
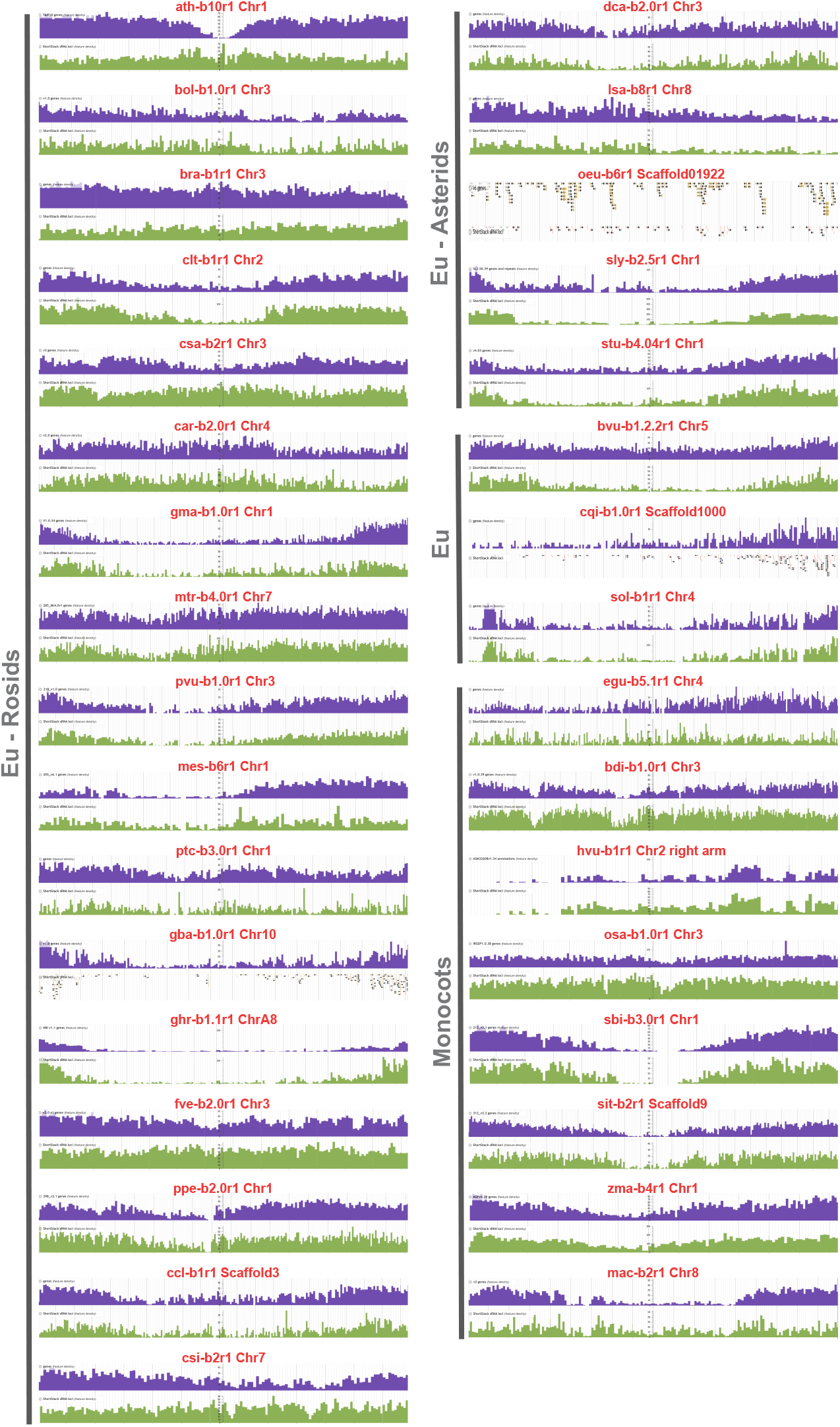
Chromosomal distribution of genes and sRNA loci. Species with a number of annotated sRNA loci high enough to have a continuous chromosomal distribution are shown, instead, species with a low number of annotated sRNA loci that are scattered across the chromosomes are not shown. For each species one representative chromosome is shown. Purple: distribution of genes. Green: distribution of sRNA loci, including the category ‘OtherRNA’, which represents a minor contribution on the total loci for the species reported here. See Table 1 for species codes. Eu: eudicots.

**Supplemental Fig. S6.**
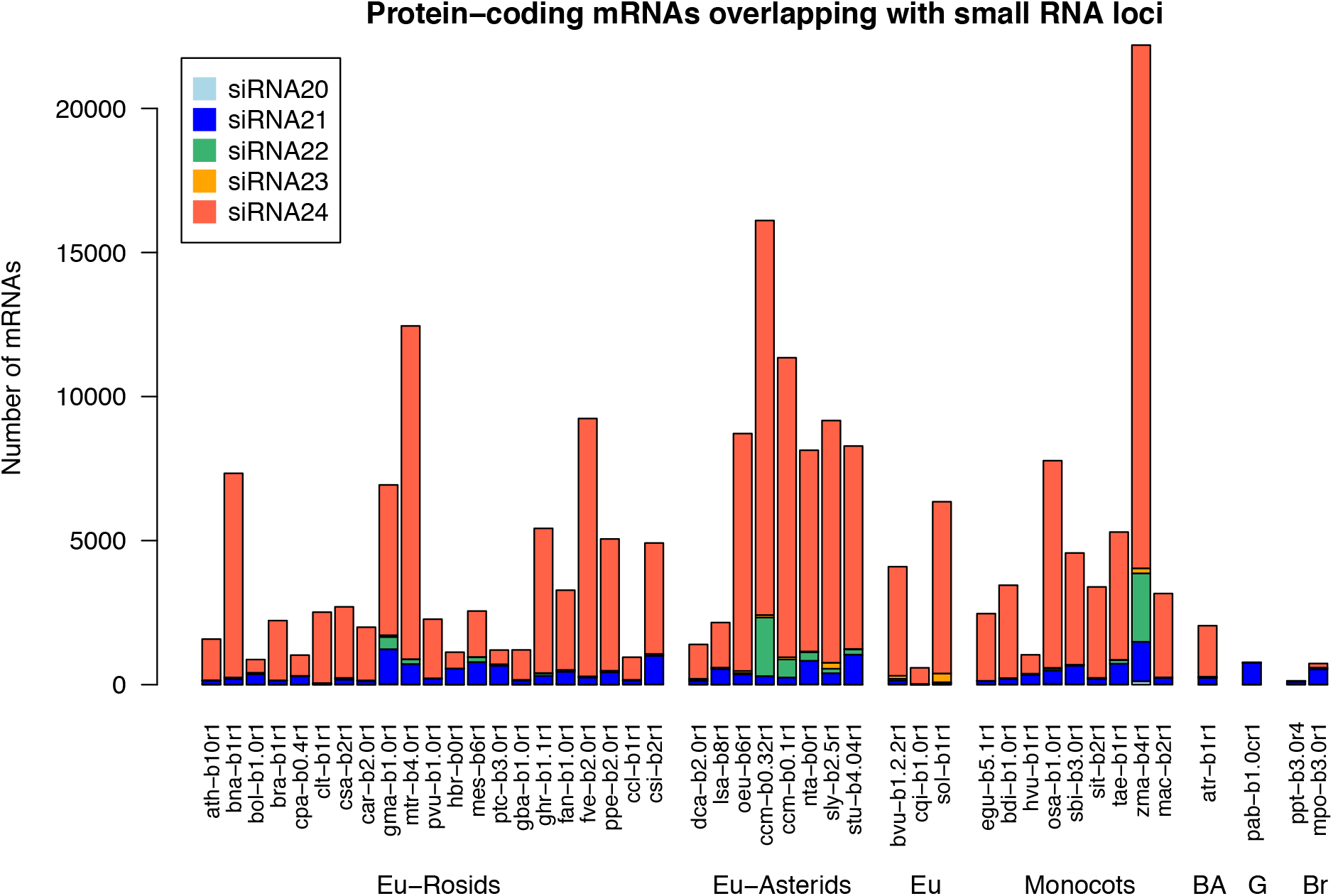
Number of protein-coding mRNAs that overlap with sRNA loci. Counts of mRNAs, containing at least one intron, that overlap with a sRNA locus for at least 25% of the length of the sRNA locus. In case of overlapping with multiple sRNA loci of different categories, the mRNA intersection was classified based on the longest overlapping sRNA locus. See Table 1 for species codes. Eu: eudicots, BA: basal angiosperm, G: gymnosperm, Br: bryophyte.

